# A unified neural representation model for spatial and semantic computations

**DOI:** 10.1101/2023.05.11.540307

**Authors:** Tatsuya Haga, Yohei Oseki, Tomoki Fukai

## Abstract

Hippocampus and entorhinal cortex encode spaces by spatially local and hexagonal grid activity patterns (place cells and grid cells), respectively. In addition, the same brain regions also implicate neural representations for non-spatial, semantic concepts (concept cells). These observations suggest that neurocomputational mechanisms for spatial knowledge and semantic concepts are related in the brain. However, the exact relationship remains to be understood. Here we show a mathematical correspondence between a value function for goal-directed spatial navigation and an information measure for word embedding models in natural language processing. Based on this relationship, we integrate spatial and semantic computations into a neural representation model called as “disentangled successor information” (DSI). DSI generates biologically plausible neural representations: spatial representations like place cells and grid cells, and concept-specific word representations which resemble concept cells. Furthermore, with DSI representations, we can perform inferences of spatial contexts and words by a common computational framework based on simple arithmetic operations. This computation can be biologically interpreted by partial modulations of cell assemblies of non-grid cells and concept cells. Our model offers a theoretical connection of spatial and semantic computations and suggests possible computational roles of hippocampal and entorhinal neural representations.

## Introduction

In the brain, place cells in the hippocampus (HPC) and grid cells in the entorhinal cortex (EC) represent spaces by spatially local and hexagonal grid activity patterns, respectively (1–4). Place cells can directly support spatial recognition and memory, whereas grid cells are considered as the basis of robust spatial navigation and path integration. Theoretically, an animal can estimate the direction to a goal when grid representations of a current position and a goal position are given (vector-based spatial navigation) (5). Furthermore, self-position can be estimated by integrating self-motions when sensory information is not available (path integration) (6, 7). These functions are bases of robust spatial navigation by animals. In addition, recent experiments suggest that grid representations in EC appear not only for physical space but also for 2-dimensional perceptual or conceptual space (e.g. 2-D visual (8), olfactory (9), social (10) spaces, and pseudowords (11, 12) and objects(13) associated to 2-D structures), and those representations are the basis of vector-based conceptual inference. HPC also exhibits neural representations for conceptual spaces (14, 15).

Because spatial navigation is a special case of decision making, we can hypothesize that such spatial representations in HPC and EC are designed for efficient reinforcement learning. Previous computational studies suggest that successor representation (SR), which was originally proposed for efficient evaluation of value functions (16, 17), can explain various experimental observations of place cells and grid cells(18, 19). Further experiments confirmed the existence of SR-like representations in the brain (19–21). Furthermore, default representations (DR) based on the theory of linear reinforcement learning (22, 23) explains that spatial representations in EC are related to flexible behaviors (24). These theories provide the formal explanation of the design principle of spatial representations like place cells and grid cells.

However, in HPC and EC, there are also neurons representing non-spatial semantic concepts which are called as “concept cells” (25–27). Concept cells respond to specific concepts, namely, stimuli related to a specific person, a famous place, or a specific category like “foods” and “clothes”, irrespectively of sensory modality (images, written or spoken words). Those cells are activated not only by external stimuli but also by imagery (28) and memory recall (29), suggesting that their activities are not merely sensory responses but conceptual representations of the high-level cognitive process. Because these cells have been found in HPC and EC, it is possible that concept cells can also be understood from the theory of spatial representations. However, how the formal spatial representation model like SR can be extended to such conceptual representations has not been fully understood.

In this paper, we show mathematical correspondence between reinforcement learning and natural language processing (NLP), from which we derive a unified neural representation model called “disentangled successor information” (DSI). We extend SR to a quantity called successor information (SI), which is equivalent to both a value function for spatial navigation in linear reinforcement learning (22–24) and an information measure used in word embedding models in NLP (30–33). Therefore, SI can be regarded as an integrated model of those two computational domains. DSI is representation vectors obtained from dimension reduction of SI under biologically plausible constraints such as non-negativity and decorrelation. In 2-D spaces, DSI forms place and grid representations corresponding to place cells and grid cells, supporting near-optimal decision making for spatial navigation. When we apply DSI to languages, DSI forms word representations in which each unit is activated by a specific concept like concept cells. Therefore, DSI representations can be interpreted as spatial and semantic representations in HPC and EC.

Furthermore, DSI model offers a common computational mechanism for spatial and semantic inferences. We show that DSI can perform analogical inference of words like word embedding models. Unlike conventional word embedding models, DSI enables the inference by switching only a few units, which can be biologically interpreted as a partial recombination of concept-cell assemblies. Intriguingly, we found that the same computational framework enables analogical inference of spatial contexts by combining previously learned spatial representations. As was suggested in experiments (34), non-grid spatial representations rather than grid representations are crucial for this computation of spatial contexts. Thus, our model provides a shared computational framework behind spatial and semantic inferences as well as a correspondence between non-grid cells and concept cells in biological computation.

Previous computational studies have revealed how spatial (18, 20, 24, 35–40) and conceptual (41–45) representations emerge and function in the brain. However, theoretical understanding of the relationship between spatial and conceptual representations has been scarce, especially for highly complex conceptual structures like languages. Although several models have recently worked on this problem (46–49), it is often difficult to discuss computational properties of neural representations other than place cells and grid cells, such as concept cells (25–27) and non-grid spatial representations that represent contextual information in EC (34, 50). Our model reveals a strong theoretical connection between spatial and semantic computations in the brain, which are biologically interpretable at unit-level and at population-level (Figure 1). From another perspective, our model suggests that word embedding models in NLP can be grounded on processing in the human brain.

**Figure 1.**
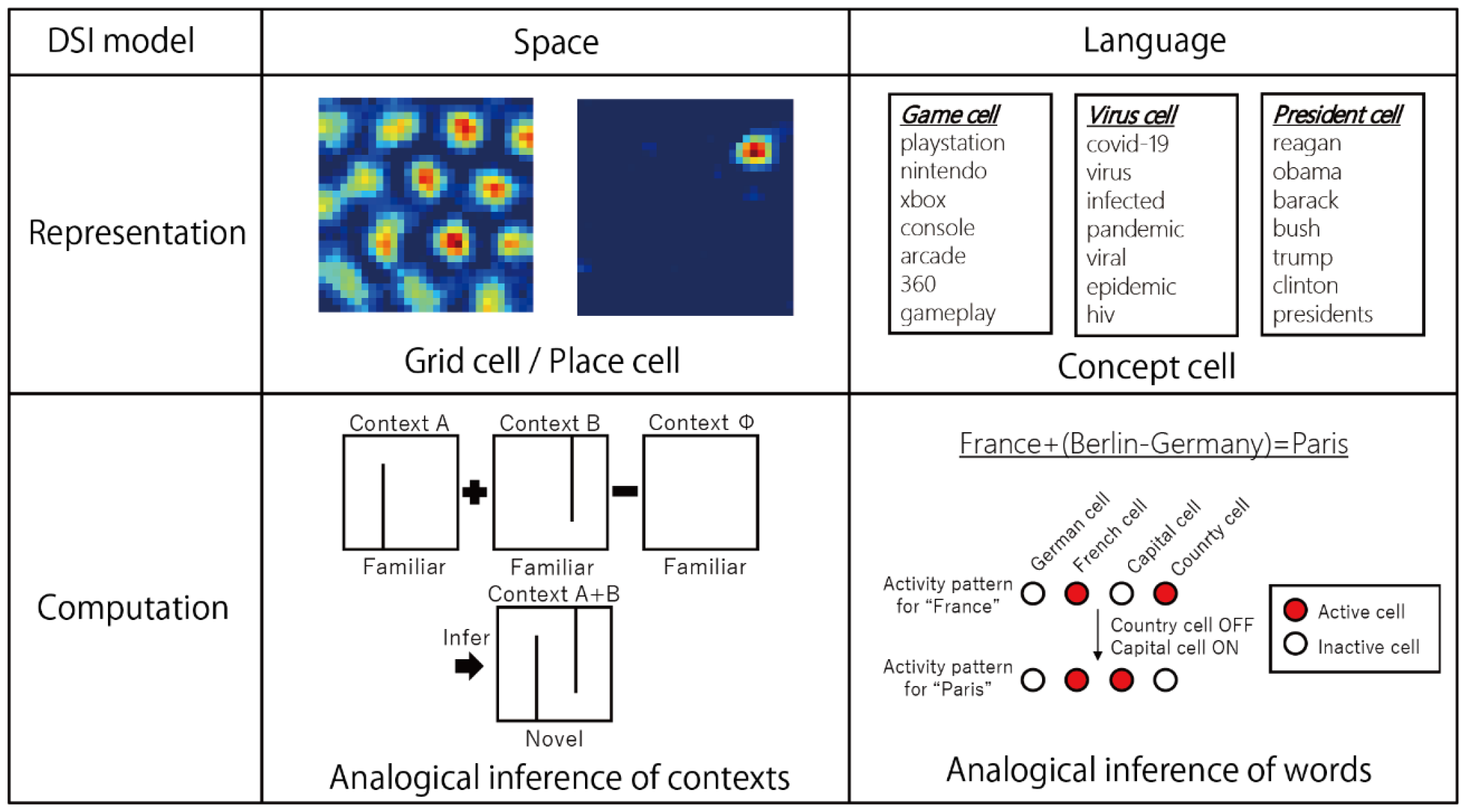
Summary of DSI model. DSI creates spatial representations like grid cells or place cells, whereas DSI also creates concept-specific word representations like concept cells. Furthermore, DSI offers a common computational framework for spatial and semantic inferences based on analogical relationships.

## Successor information

We introduce a quantity called as SI as an extension of SR (16, 18, 19). Let us assume *N*_*s*_ discrete states (positions) exist in the environment. SR between two states *s* and *s*^′^ is defined as

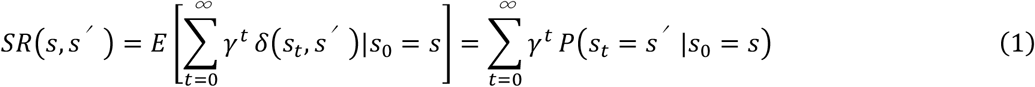

where *δ*(*i,j*) is Kronecker’s delta and *γ* is a discount factor. Based on SR, we define successor information (SI) and positive successor information (PSI) as follows.

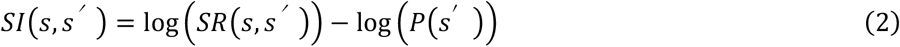

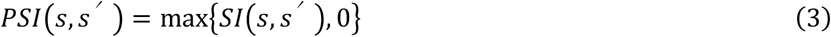

Intuitively, log (*SR*(*s,s’*)) measures temporal proximity (reachability) between states, and self-information − log (*P*(*s*^′^)) normalizes log (*SR*(*s,s’*)) that tends to increases as a function of the occurrence frequency of the state. PSI neglects weak relationships between distant states by rectifying SI. We determined this mathematical form of SI so that it corresponds to both a value function for goal-directed spatial navigation and an information measure for word embedding models. This indicates that there is a mathematical correspondence between the reinforcement learning and word embedding models.

First, SI corresponds to a value function of linear reinforcement learning (22–24) in a specific setting of spatial navigation. Linear reinforcement learning assumes default policy and imposes additional penalty on deviation from default policy, then we can obtain value functions explicitly by solving linear equations. Let us consider a specific condition in which the environment consists of non-terminal states, and a virtual terminal state is attached to a goal state *s*_*G*_ arbitrarily chosen from non-terminal states. When the agent gets to the goal, it transits to the terminal state and obtain positive reward. Furthermore, we assume that rewards at non-terminal states are uniformly negative so that the agent has to take a short path to goal to maximize reward. In this setting for goal-directed spatial navigation, we can obtain value functions *v*^*\**^(*s*) in linear reinforcement learning as

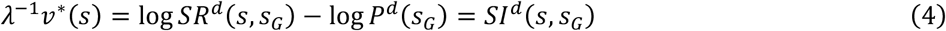

where *SR* ^*d*^(*s, s*_*G*_) and *SI* ^*d*^(*s, s*_*G*_) are SR and SI under the default policy, respectively, *P*^*d*^(*s*_*G*_) is a probability of visiting the state *s*_*G*_ under the default policy, and *λ* is a parameter representing the relative weight of penalty on deviation from default policy (see Methods “Mathematical relationship of SI and reinforcement learning” for details of derivation). Therefore, SI is proportional to value functions for spatial navigation.

Second, DSI is related to word embedding models in NLP (30–33, 51). In linguistics, pointwise mutual information (PMI) and positive pointwise mutual information (PPMI) are used to measure the degree of coincidence between two words(33). They are defined as

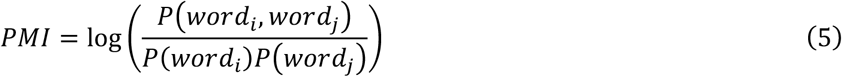

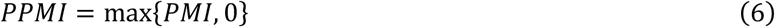

where *P*(*word*_*i*_, *word*_*j*_)is a coincidence probability of two words (in a certain temporal window). It has been proven that dimension reduction of PMI approximates a conventional word embedding model skip-gram(30, 31), and similar performance is obtained using PPMI (33). SI can be written as

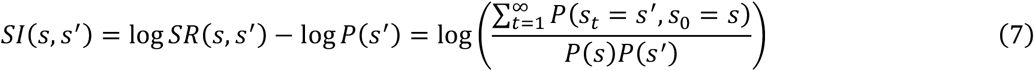

In this formulation, we can see mathematical correspondence between PMI and SI by regarding words as states (*s* = *word*_*i*_, *s*^′^ = *word*_*j*_), thus the correspondence between PPMI and PSI. The difference is how to count coincidence: the coincidence in SI is evaluated with an asymmetric exponential kernel as in SR, in contrast that a symmetric rectangular temporal window is often used in typical word embedding. Because of this relationship, we can expect that representation vectors obtained by dimension reduction of a PSI matrix has similar properties to word embedding models.

### Disentangled successor information

We perform the dimension reduction of the PSI matrix using extensions of nonnegative matrix factorization (NMF) (52) with additional constraints. Then, we finally obtain *D*-dimensional non-negative vectors 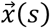 and 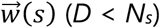 which satisfies 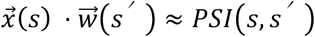. We call those representation vectors as disentangled successor information (DSI). These representation vectors correspond to those obtained in word embedding models because of the relationship between PSI and PMI. Furthermore, from the mathematical correspondence shown in the previous section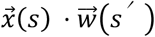 approximates a value function of *s* when *s*^*′*^ is given as a goal. Therefore, we basically regard 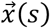 as a representation of each state, and 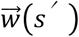 represents a temporary goal. Below, we call 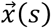 as a DSI representation vector unless otherwise specified.

In this study, we test two types of constraints for dimension reduction, and we call the resultant vectors as DSI-decorr or DSI-sparse depending on settings. The first one, DSI-decorr was constrained by non-negativity, decorrelation, and L-2 regularization. Previous theoretical studies have shown that these constraints are important for generation of grid representations (35–40, 53, 54). The other one, DSI-sparse was obtained under non-negativity and L-1 sparse constraint. A previous study in word embedding suggests that these constraints enable generation of word representation vectors with conceptual specificity (51). Therefore, we can expect that either or both of those constraints achieves simultaneous generation of spatial representations and semantic representations which correspond to grid cells and concept cells, respectively. These constraints are biologically plausible because neural activities are basically non-negative, and decorrelation and sparsification is possible through lateral inhibition. For details of dimension reduction, see Methods “Details of dimension reduction for DSI vectors”.

### Emergence of spatial representations like grid cells and place cells

Below, we demonstrate properties of DSI representations through simulations. First, we checked spatial representations that the DSI model forms in a 2-D space. As an environment, we assumed a square room tiled with 30 × 30 discrete states (Figure 2A). We generated sequences of states by random walks in the room, from which we calculated 100-dimensional DSI representation vectors for states (places) (see Methods “Learning DSI in 2-D spaces” for the detail of the simulation).

**Figure 2.**
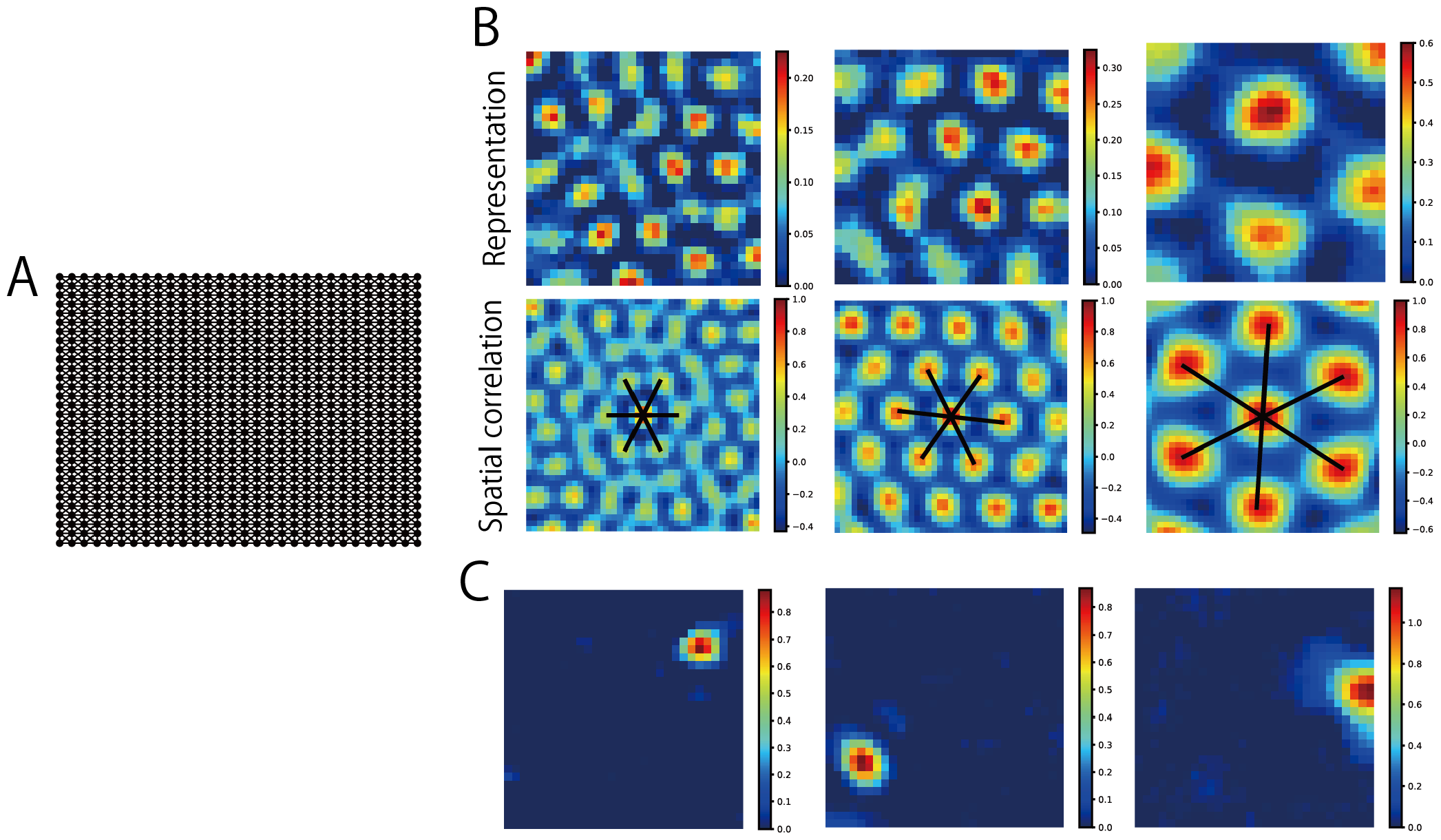
DSI representations for the space. (A) The square room tiled with 30×30 discrete states. (B) Grid-like spatial representations generated by DSI-decorr (upper) and their spatial autocorrelation (lower). (C) Spatial representations like place cells generated by DSI-sparse.

Here we call each dimension of DSI representation vectors 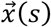 as a neural “unit”, and we regard a value in each dimension at each state as a neural activity (or a neural representation). Then, in DSI-decorr vectors, many units exhibited grid activity patterns in the space (Figure 2B). We performed a gridness analysis that has been used in previous studies (37, 55–57) and found that 27.6±5.6 % of units (average±std of 5 simulations with different random seeds) were classified as grid cells. Similarly, 30.6±4.0 % of units in *w*(*s*) were classified as grid cells. On the other hand, DSI-sparse generated spatially local representations which resemble hippocampal place cells (Figure 2D). Therefore, we regard DSI-decorr and DSI-sparse as models of EC and HPC representations, respectively. Below, we mainly show results of DSI-decorr but DSI-sparse also gave qualitatively similar performances in most cases.

### Path integration and spatial navigation by DSI representations

If DSI representations are a plausible model of place cells and grid cells, they should support path integration and spatial navigation, which are important functions of HPC and EC. First, we performed path integration using DSI representations. In the path integration task, after a starting location (state) is given, an agent has to estimate the current position of itself by integrating only self-movement information. To solve the task, we estimated spatial representations by movement-conditional recurrent weights at each time step (see Methods “Path integration by DSI vectors” for details). This strategy has been used in previous studies such as grid cell modeling (39) and action-conditional video prediction (58). This mechanism is also consistent with a conventional biological model for path integration in which head direction signals activate one of attractor networks specialized for different directional shifts of grid patterns (6, 7, 36). As shown in Figure 3A (DSI-decorr) and Supplementary Figure 1 (DSI-sparse), this strategy gave accurate estimation of the spatial path from movement signals. To quantify the accuracy, we evaluated the success rate of estimation of spatial paths generated from random starting points and sequences of ten random movements. Estimation was correct in 969.4±8.3 trials (DSI-decorr) and 759.2±14.4 trials (DSI-sparse) out of 1,000 trials (average±std of 5 simulations with different random seeds), which suggests that path integration was highly accurate especially with DSI-decorr.

**Figure 3.**
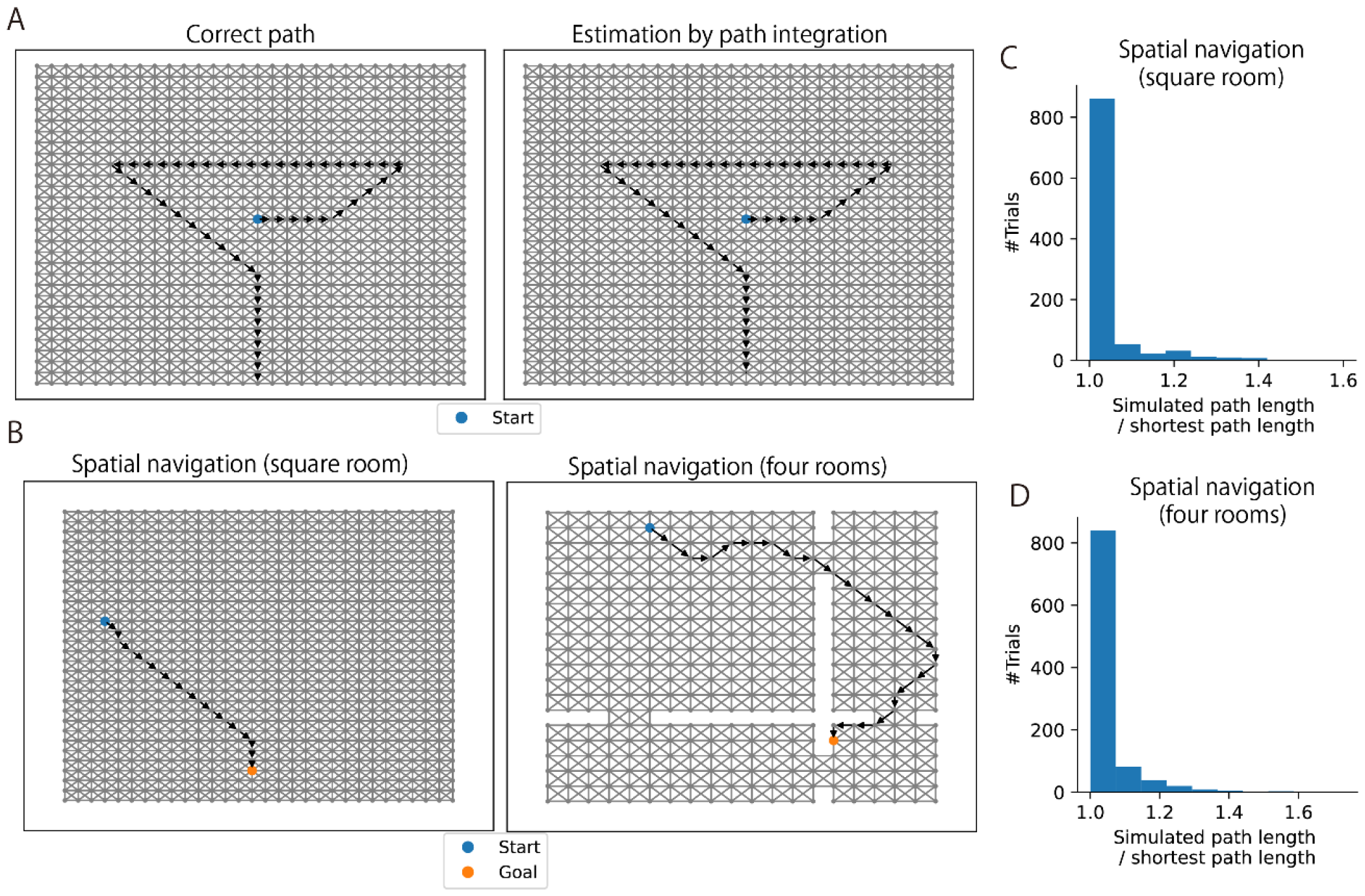
Spatial navigation using DSI representation vectors (DSI-decorr). (A) Path integration using DSI model. (Left) Actual path. (Right) Path estimated from DSI vectors updated by movement information. (B) Example spatial paths obtained by DSI-based navigation. (C) A histogram of path lengths in 1,000 trials of spatial navigation in the square-room environment. Note that a start and a goal were randomly determined in each trial, and we normalized a simulated path length by the shortest path length between the start and the goal. (D) A histogram of path lengths in the four-room environment.

Next, we tested spatial navigation by DSI representations. We assume that a start location and a goal location are randomly given in each trial and an agent has to navigate between them. To solve the task, we defined a vector-based state transition rule that approximates value-based decision making based on the relationship between DSI and value functions (see Methods “Goal-directed spatial navigation by DSI vectors” for details). Because of the constraints in the model and the approximation error, this rule did not always give optimal navigation (the shortest path from the start to the goal). However, the agent could take relatively short paths (Figure 3B, left and Supplementary Figure 1) and simulated path lengths were close to the shortest path between the start and the goal (Figure 3C and Supplementary Figure 2). Furthermore, the agent could also perform near-optimal spatial navigation in a structure with separated and interconnected rooms (Figure 3B, right, Figure 3D, and Supplementary Figure 1). These results suggest that DSI representations can support efficient spatial navigation.

### Emergence of concept-specific representations for words

Next, we show that the same DSI model can learn conceptual representations from linguistic inputs. We used text data taken from English Wikipedia, which contains 124M tokens and 9376 words (see Methods “Learning DSI from text data” for the detail of preprocessing). To construct DSI representations, we regarded each word as a state, and considered the text data as a sequence of 9376 states (*N*_*s*_ = 9376). Then, we applied the same learning procedure as in the experiment of 2-D spaces. We obtained 300-dimensional DSI representation vectors for each word.

As in the case of spatial representations, we regard each dimension of representation vectors as a neural unit, and checked how various words activate those units. Specifically, we listed ten words that elicited the highest activities in each unit (TOP-10 words). Then, we found that many units are activated by words related to specific concepts (Figure 4A; other examples in Supplementary Figure 3), which could be named as “game cell” or “president cell”, for example. We quantified this conceptual specificity through WordNet-based semantic similarity between words (59). We compared mean similarity among TOP-10 words and a null distribution of similarity between random word pairs, by which we determined statistically significant concept-specific units and quantified the degree of conceptual specificity of each unit (see Methods “Quantitative evaluation of conceptual specificity” for details). As shown in Figure 4B and 4C, both DSI-decorr and DSI-sparse exhibited the larger number of concept-specific units and higher average conceptual specificity than other well-established word embedding models such as skip-gram and GloVe (30–33). We also analyzed conceptual specificity of representations in the embedding layer of pretrained BERT model(60, 61), which was lower than DSI. This result shows that DSI model forms more concept-specific representations than typical word embedding models in NLP. Remarkably, removal of the non-negativity constraint from DSI-decorr significantly decreased conceptual specificity. This result suggests that non-negativity is an essential constraint for concept-specific word representations as well as hexagonal grid spatial representations (35, 36, 53), and sparsity is not necessary as proposed in the previous study (51). It is also notable that continuous bag-of-words (CBOW) model (31) showed almost same conceptual specificity with DSI. However, we see a functional difference between concept-specific representations in DSI and CBOW later.

**Figure 4.**
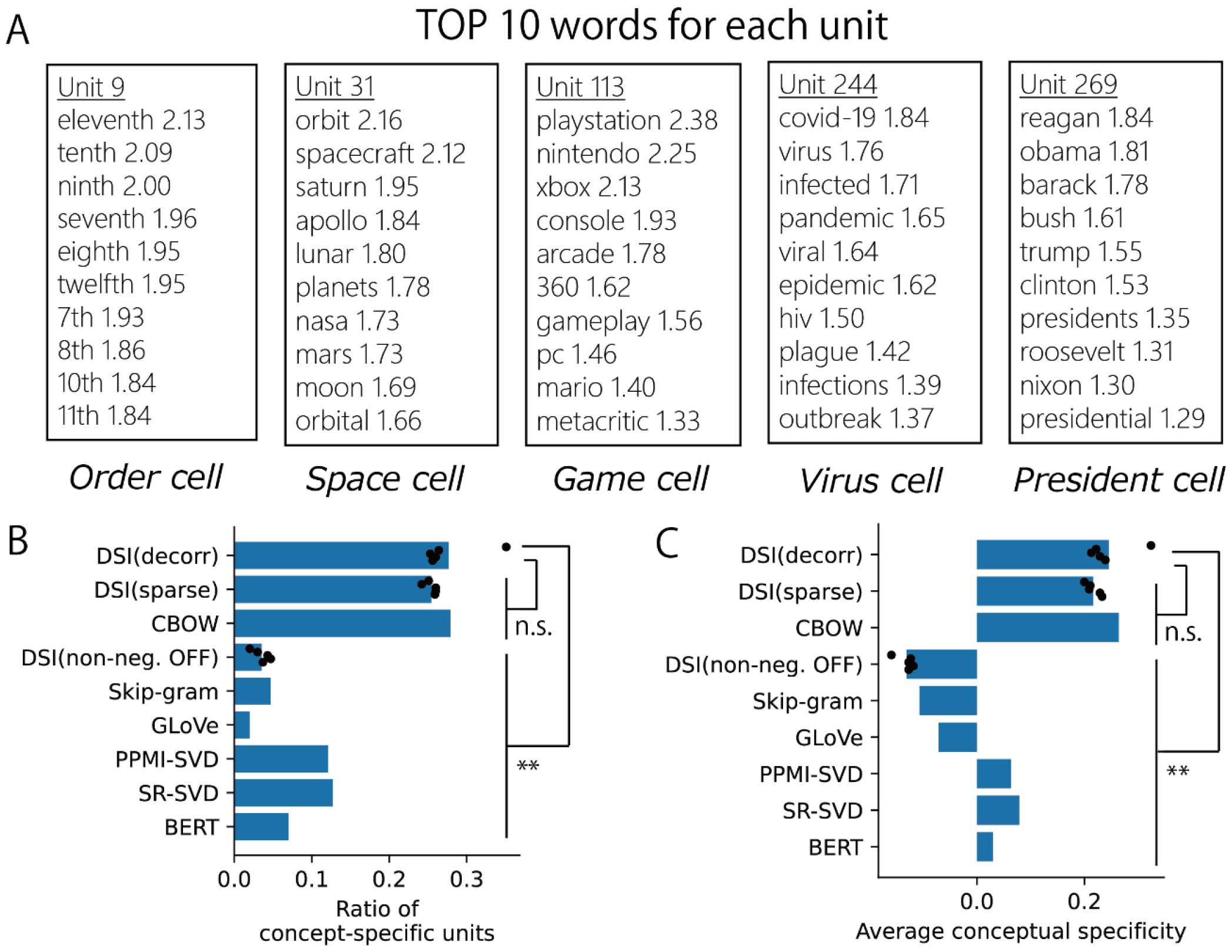
Concept-specific representations formed by DSI-decorr. (A) Ten words that gave the highest activation (TOP-10 words) are shown. We also marked each unit with a descriptive label. (B) Ratio of concept-specific units (dimensions) in word representation vectors obtained by various methods. For DSI, dots indicate 5 trials with different random seeds (different initial values for learning); bars indicate means of those 5 simulations. We compared DSI-decorr and other DSI models by 2-sample t-tests, and DSI-decorr and other word embedding methods by 1-sample t-tests. (C) Average conceptual specificity of all units (dimensions) in word representation vectors obtained by various methods. We performed same statistical analyses with (B). **P<0.01; n.s., not significant. All statistical tests were two-sided t-tests and significance thresholds were modified by Bonferroni correction. Details of statistical analyses are shown in Supplementary Table 1.

### DSI representation vectors capture the semantic structure of words

Concept cells are considered to represent semantic relationships between concepts at the level of population activity patterns (25). Correspondingly, DSI vectors also represent the semantic structure of words because representation vectors of word embedding models have such property (30–33). We evaluated how word similarity is captured by cosine similarity of DSI vectors using the evaluation procedure in NLP (30–33). We calculated cosine similarity between representation vectors of word pairs and evaluated the rank correlation between those cosine similarities and human word similarities (WS353 dataset (62); 248/345 word pairs were used). As a result, DSI showed high correlations that are comparable to word embedding models (Figure 5A). This indicates that DSI captures the semantic structure of words at the level of population activity patterns.

**Figure 5.**
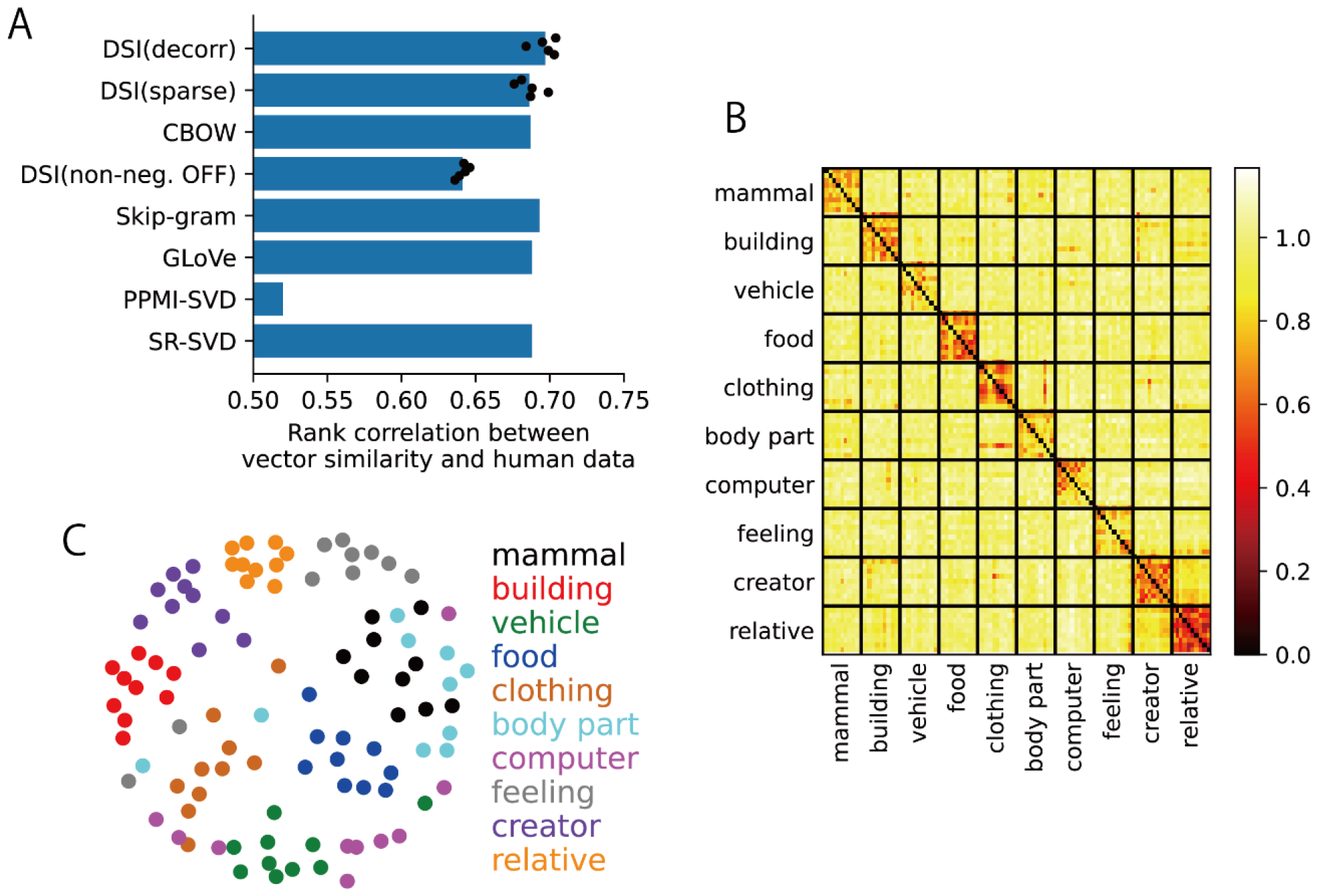
DSI representations capture the semantic structure of words at the population level (DSI-decorr). (A) The rank correlation of word similarity evaluated by word representation vectors (cosine similarity) and humans (WS353 dataset). For DSI, dots indicate 5 trials with different random seeds (different initial values for learning); bars indicate means of those 5 simulations. (B) Dissimilarity matrix between DSI representation vectors for 100 words in 10 semantic categories (DSI-decorr). We selected 10 words in each category. We used same dissimilarity metric with Reber et al. (2019) (1 - Pearson’s correlation coefficient). (C) Visualization of the representational structure of DSI using MDS based on the dissimilarity matrix shown in (B). Each color corresponds to a semantic category.

This property suggests that the representational structure of DSI vectors is consistent with the experimental observation that population-level pattern similarity of concept cell activities represents semantic categories (26). We considered 10 semantic categories and chose 10 words corresponding to each category (see Methods “Evaluation of the semantic structure of DSI vectors” for the choice of words) and evaluated dissimilarity between DSI representation vectors of those 100 words using the same metric with the experimental work (26) (1 - Pearson’s correlation coefficient). Dissimilarity matrix in Figure 5B and Supplementary Figure 4 clearly shows that representations were similar within each semantic category. In the visualization of the representational structure by multidimensional scaling (MDS), we can also see clustering of words corresponding to 10 semantic categories (Figure 5C and Supplementary Figure 4).

### Analogical inference by partial recombination of assemblies of conceptual units

So far, we have analyzed how DSI represents spaces and semantic concepts. In this section, we discuss how DSI can support computation of semantic concepts. Specifically, we argue that DSI provides a biological interpretable mechanism for inference of words. Previous studies in NLP (30, 31) have shown that arithmetic calculation of word representation vectors enables inference of words based on analogical relationships. For example, when we consider an analogical relationship “man is to woman as king is to queen”, the calculation of word representation vectors as 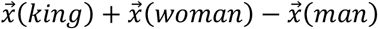 gives a similar vector to 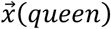. For this analogical inference task, DSI vectors show comparable performance with conventional word embedding models (evaluation by Mikolov’s dataset (30, 31); 3157/19544 questions were used; see Methods “Analogical inference of words” for details) (Figure 6D, success rates at “300 (Full)”). However, unlike the conventional word embedding, each word is represented by a combination of concept-specific units (a population activity pattern of concept cells) in DSI. In this case, we can interpret its analogical inference as a partial recombination of those assemblies to create novel word representations. Below, we give an intuitive explanation of this property of DSI.

**Figure 6.**
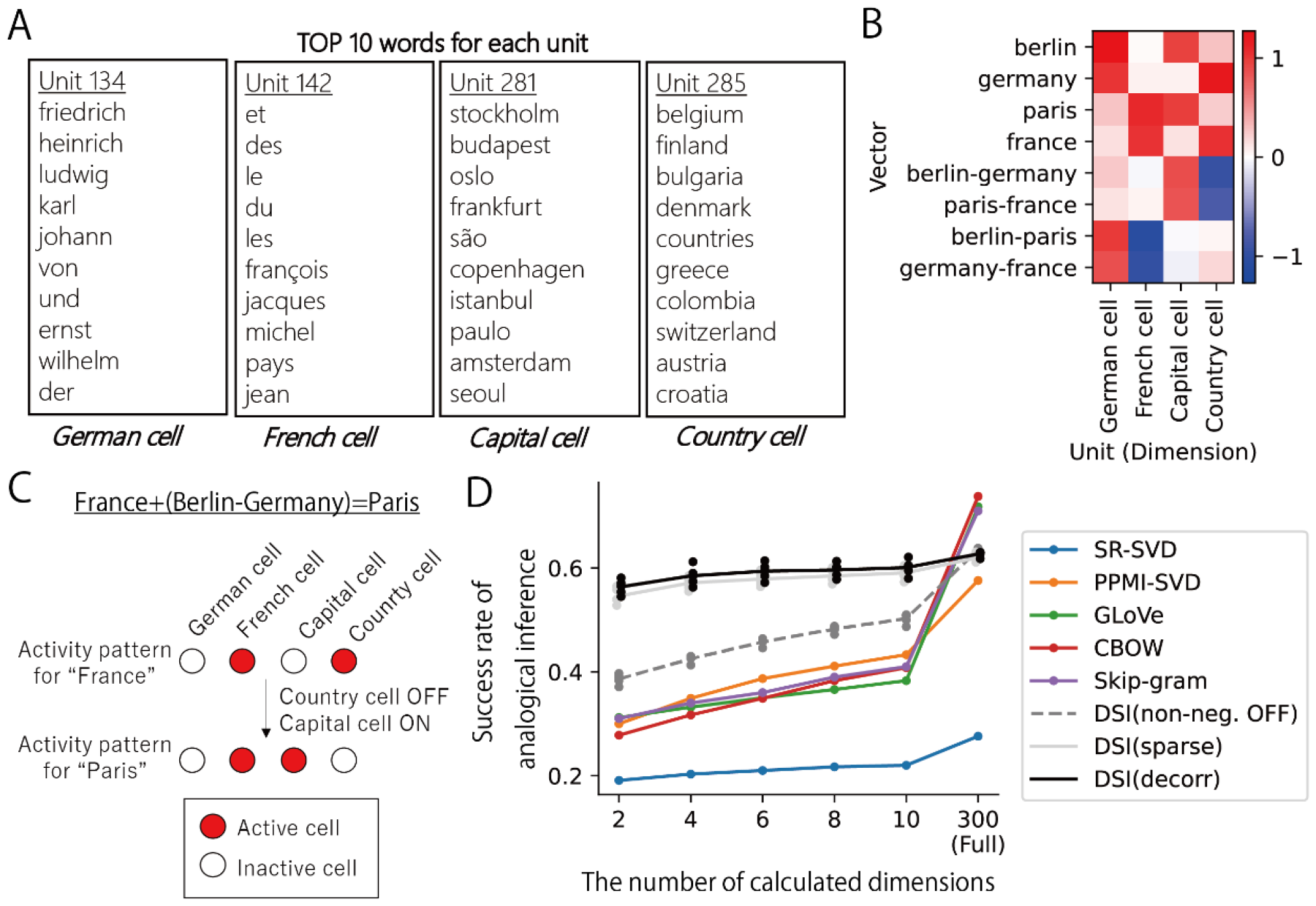
Analogical inference as a sparse recombination of conceptual units. (A) TOP-10 words of four representative units obtained by DSI-decorr and their interpretation. (B) Values of four representative units in DSI vectors for France, Germany, Paris, Berlin, and difference vectors between them. (C) Interpretation of analogical inference as sparse recombination of concept cells. (D) Success rates of analogical inference task (Mikolov’s dataset) by calculating the limited number of dimensions in word representation vectors. For DSI, dots indicate 5 trials with different random seeds (different initial values for learning); lines indicate means of those 5 simulations. Success rates of DSI-decorr were not significantly different from DSI-sparse (P>0.05, 2-sample t-test) but higher than all other methods in the condition of 2, 4, 6, 8, and 10 dimensions. (P<0.01, 2-sample t-test for DSI (non-neg. OFF) and 1-sample t-tests for other word embedding methods). All statistical tests were two-sided t-tests and significance thresholds were modified by Bonferroni correction. Details of statistical analyses are shown in Supplementary Table 2.

First, we analyzed the ratio of each element to the sum of all elements in DSI vectors. We found that even the largest element accounted for 5% of the sum of all elements on average in the case of DSI-decorr (Supplementary Figure 5). This implies that DSI vectors for words are non-sparse and distributed; thus, each word is represented by a combination of multiple conceptual units.

Next, we inspected representations of a set of words as an example: France, Paris, Germany and Berlin. In those words, we can see two analogical relationships (country-capital and French-German relationships). We identified the most active units in DSI vectors for those words and listed the TOP-10 words for identified units. We could find four representative units that we could name as “German cell”, “French cell”, “country cell” and “capital cell” (Figure 6A). We could see that 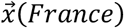 is represented by the combination of the French cell and the country cell, whereas 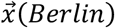 is represented by the combination of the German cell and the capital cell, and so on (Figure 6B). This example gives a simple interpretation of word similarity in the DSI vector space discussed in the previous section. If words are similar, they share a large number of active conceptual units, like the country cell shared by representations of France and Germany. Thus, semantic similarity between words increases cosine similarity between word vectors.

Then, the difference vectors between words correspond to switching on and off conceptual units. For example, summing the difference vector of 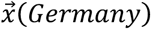 and 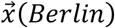 can turn on the capital cell and turn off the country cell (Figure 6B). Similarly, transformation from 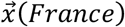 to 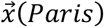 corresponds to the activation of the capital cell and deactivation of the country cell, enabling this transformation by summing the difference of 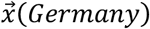 and 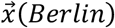. Thus, we can biologically interpret the procedure of analogical inference as a partial recombination of an assembly of concept cells (Figure 6C). If neural circuits in HPC and EC have learned transformation of 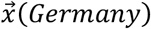 to 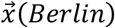 by sparse switching of concept cells, the same operation enables 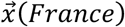 to 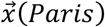. Here, the mathematical mechanism of the analogical inference is the same as the conventional word embedding models. However, a unique feature of our DSI representations is that those analogical relationships are factorized into separated units, making the modulation of a few units enough to perform the inference. We speculate that constraints in our dimension reduction method (especially non-negativity) are sufficient conditions to align each semantic factor to each axis of the vector space.

Next, we check whether the property of DSI mentioned above is general across a wide repertoire of words. If each analogical relationship is actually confined to a specific dimension, switching on and off only a few dimensions is enough for the inference, and the calculation of a whole vector is unnecessary. We tested this hypothesis by an analogical inference task using Mikolov’s dataset, limiting the number of calculated dimensions. Namely, when we calculated 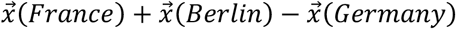 to obtain 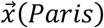, we identified a few dimensions that had maximum and minimum values in 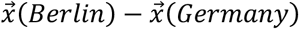 (that are “country cell” and “capital cell”). We summed only those dimensions to 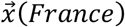. This partial calculation of vectors did not degrade the overall task performance of DSI-decorr and DSI-sparse (even 2 out of 300 dimensions did not largely alter the performance) (Figure 6D). In contrast, the performance of other word embedding models was significantly impaired (Figure 6D). This result suggests that the partial recombination of concept-specific units enabled the analogical inference of various words, and this property is unique to DSI model. Removal of the non-negativity constraint significantly impaired the performance of DSI-decorr, which further suggests the importance of non-negativity (Figure 6D). Notably, CBOW model, which showed high conceptual specificity (Figure 3), did not work properly in this inference task with partial calculations (Figure 6D). This suggests that apparent conceptual specificity is not a sufficient condition for such calculation.

### Analogical inference of spatial contexts by modulating non-grid spatial representations

In the previous section, we discussed analogical inference of words. We found that we can apply the same computational framework to spatial representations. Furthermore, our results suggest a novel computational role of non-grid (heterogenous) spatial representations that have been experimentally found in EC (34, 50).

First, we constructed DSI representation vectors (DSI-decorr) for spatial contexts A, B, and Φ, which have different barrier layouts (Figure 7A, Supplementary Figure 6). Then, we created composite representation vectors for a novel context A+B by simply combining the spatial representation vectors for familiar contexts A, B, and Φ as “A+B-Φ” (Figure 7A, see Methods “Analogical inference of spatial contexts” for details). We devised this computation based on the analogical inference of words: the relationship of Φ to B corresponds to the relationship of A to A+B (whether a barrier exists at a specific position or not). We tested goal-directed spatial navigation (as described in the previous section) in three spatial contexts A, B, and A+B, by using the learned representations for A, B, and Φ, and those generated for A+B. Naturally, the representation vectors for A and B gave the best performance in contexts A and B, respectively (Figure 7B and 7C). Remarkably, the composite representation vector “A+B-Φ” achieved the best performance in the context A+B (Figure 7B and 7C). This result corresponds to the vector computation for word inference, suggesting that the computational framework for semantic concepts is also valid for inferring the spatial contexts. We confirmed that this inference also worked in another spatial setting (Supplementary Figure 7). However, the inference did not improve spatial navigation when two barriers were spatially connected in the context A+B (Supplementary Figure 8). We speculate that this simple computational scheme works only if the interaction between barriers is small. Therefore, we conducted further analyses below in the settings used in Figure 7 and Supplementary Figure 7.

**Figure 7.**
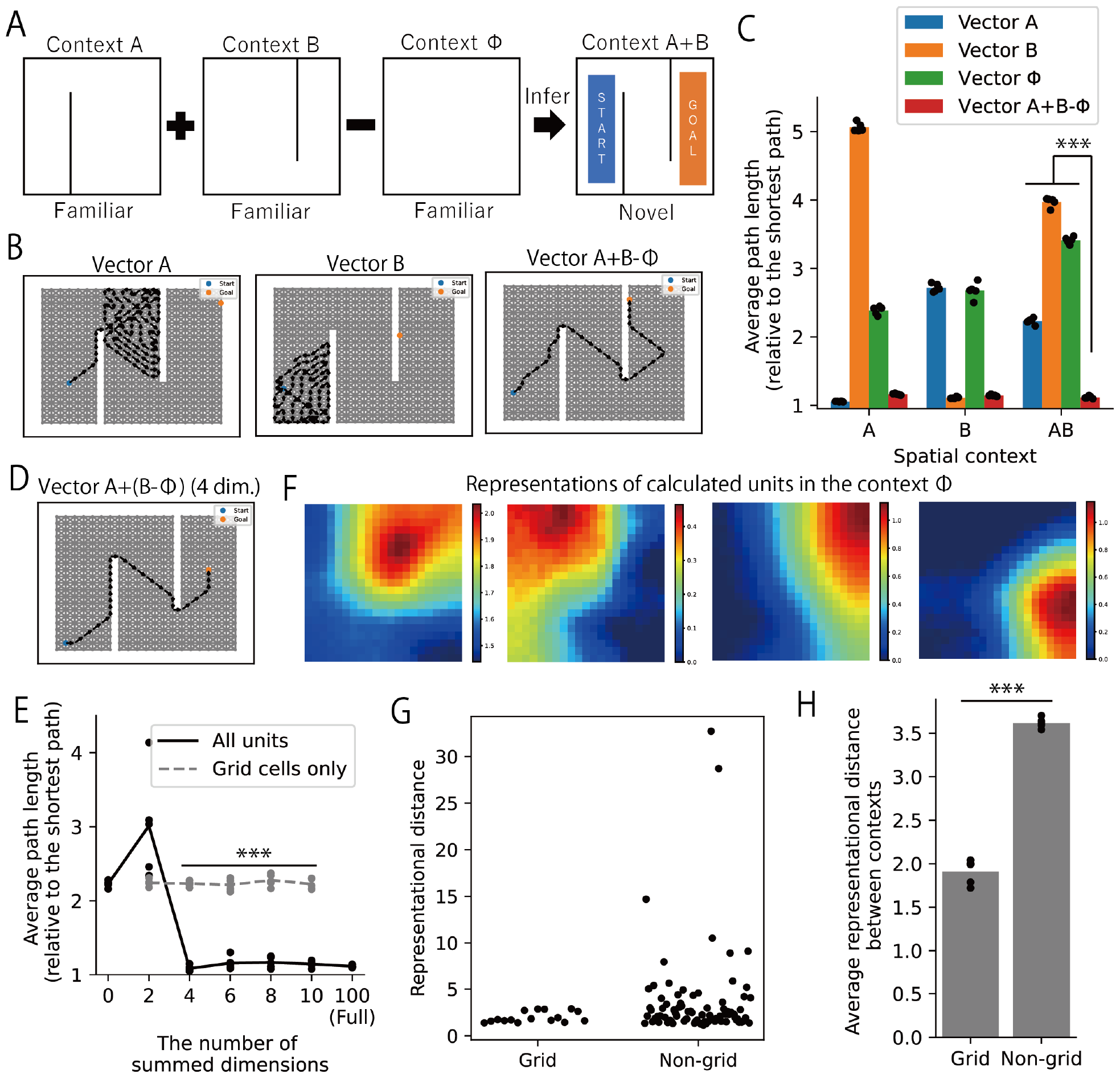
Composite spatial representations enable navigation in a novel spatial context. (A) We constructed representation vectors for a novel context A+B by arithmetic composition of DSI representation vectors for three familiar contexts A, B, and Φ. The start and the goal in each navigation trial were randomly positioned in the colored area. (B) Example spatial paths by spatial navigation using representation vectors learned in the context A (left), B (middle), and composite representation vectors A+B-Φ (right). (C) Average path lengths in 1,000 trials of spatial navigation under various settings of representation vectors and contexts. Note that we normalized a path length by the shortest path length between the start and the goal in each trial. Dots indicate 5 simulations with different random seeds (different initial values for learning and simulations); bars indicate means of those 5 simulations. (D) An example of spatial navigation performed by representation vectors created by calculating only 4 dimensions of the vector A. (E) Average path lengths by the spatial navigation using the composite representation vectors in which we summed the given number of dimensions. Dots indicate 5 simulations with different random seeds (different initial values for learning and simulations); lines indicate means of those 5 simulations. 2-sample t-tests were performed between the condition in which we calculated only grid cells and the condition in which we calculated all units. (F) Spatial representations of four calculated units in the context Φ in an example of (D). Note that all these units are non-grid representations. (G) Representational distances between the context B and Φ of grid-type and non-grid-type units in a simulation. Each dot indicates a unit. (H) Average representational distances between the context B and Φ of grid-type and non-grid-type units. Dots indicate 5 simulations with different random seeds (different initial values for learning and simulations); bars indicate means of those 5 simulations. ***P<0.001. All statistical tests were two-sided t-tests and significance thresholds were modified by Bonferroni correction. Details of statistical analyses are shown in Supplementary Table 3.

We hypothesized that the relationship between spatial contexts is confined to the small number of units (dimensions) as in word representations. If this is the case, representational modulations of only a few units are enough for the inference. We identified a few units that had the largest representational differences (see Methods “Analogical inference of spatial contexts” for the definition of representational distance) between the spatial contexts B and Φ, and calculated only those units to compose representation vectors for the context A+B. We found that minimally 4 units were enough to obtain the asymptotic performance (Figure 7D and 7E). We inspected the spatial representations of these units and found that all of them were non-grid representations in the context Φ (no barriers) (Figure 7F). Consistently, we found large representational distances selectively in non-grid-type units (Figure 7G), and the average representational distance of non-gird-type units is larger than that of grid-type units (Figure 7H and Supplementary Figure 7). Strikingly, when we restricted calculation of the vectors to grid-type units (excluding non-grid-type units), the obtained composite representation vectors did not improve spatial navigation in the context A+B (Figure 7E). We obtained qualitatively same results in the other spatial setting (Supplementary Figure 7). These results suggest that information of spatial contexts (barrier settings in this case) is confined to the limited number of units that have non-grid spatial representations, and the modulation of non-grid representations is sufficient for inferring novel spatial contexts.

These results imply that the modulation of non-grid spatial representations is more informative for coding differences between spatial contexts than that of grid representations. This is consistent with the previous experimental finding that non-grid spatial representations are remapped more significantly than grid cells across different spatial contexts (34). Our model provides a novel theoretical interpretation for this experimental observation, linking the computation of spatial contexts to that of semantic concepts.

## Discussion

In this paper, we proposed a theoretically interpretable and biologically plausible neural representation model for spatial navigation and semantic concepts. Our model is mathematically related to reinforcement learning and word embedding, thus representations support spatial navigation and NLP. We demonstrated that our DSI model forms spatially local or hexagonal grid representations for the 2-D space and concept-specific representations for the linguistic inputs, which can be regarded as neural representations in HPC and EC. Finally, the same computational framework based on partial representational modulation enables the inference of words and spatial contexts, which can be biologically interpreted as the function of concept cells and non-grid cells. These results suggest that we can extend the spatial representation model of HPC and EC to learn and compute semantic concepts, which apparently seems a different computational domain from spatial navigation, in an intuitive and biologically plausible manner.

### Theory of grid and non-grid representations in EC

Our model produces grid-like representations in 2-D space, which supports path integration and spatial navigation. Previous studies have revealed that nonnegative and orthogonal constraints are important to obtain realistic grid-like representations (35, 36, 53). Furthermore, recurrent neural networks form grid-like representations through learning path integration, and those representations support efficient spatial navigation (36–40). It has also been shown that a unified model for spatial and non-spatial cognition generates grid representations (46, 47, 49). Although our model was built on the basis of those previous works, previous models have not been applied to learning of semantic concepts or other complex conceptual spaces in real-world data. Furthermore, our results revealed that the computational framework for semantic concepts (analogical inference) can be applied to spatial representations, and non-grid spatial representations were important for the inference. Our model extended the range of applicability of biologically plausible spatial representation models to semantic concepts and suggests shared computational mechanism between those two computational domains.

Furthermore, our model suggests the importance of non-grid spatial representations for inference (or switching) of spatial contexts. In EC, there are many neurons that exhibit non-grid spatial representations, and they show large remapping across contexts (34, 50). Previous models have shown that non-grid (heterogenous) spatial representations in EC support precise spatial encoding (36) and reward-based modulation of path integration (40). Our results give alternative explanation of such remapping: such remapping can support inference of navigation strategy across spatial contexts. However, it is still to be investigated whether this computational framework extends to general contextual changes such as wall colors, odors and reward settings (34, 40, 50). We speculate that this is possible if we can appropriately construct disentangled representations for general sensory and task variables (54, 63, 64).

### Implications for conceptual representations in the brain

Models of concept cells have mostly assumed sparsity and clustering to create conceptual specificity (41, 42, 46, 47, 51). In contrast, our results suggest that non-negativity is essential to create functionally useful concept-specific representations. Non-negativity is also crucial to generate biologically plausible grid representations (35, 36, 53), thus our model suggests a common constraint across spatial and conceptual representations in HPC and EC. A recent theoretical finding suggests that non-negativity and energy minimization enable disentanglement of visual, spatial, and task variables from mixed inputs (54). Our results suggest that such strategy also works in the case of semantic concepts, and it enables a simple framework for the inference in the brain.

The biological interpretation of our model predicts a population-level property of concept cells. When a human subject (or an animal) learns a novel word (or an item) that can be interpreted by the composition of previously learned concepts, the representation for the novel item can be created by partial recombination of previous formed concept cells, instead of creating novel concept cells. Concept cells exhibit concept-specific activities, but they can simultaneously work as building blocks to represent specific items as assemblies (25, 26). Thus, recombination of those assemblies is useful for creation (or prediction) of novel conceptual representations. This strategy is beneficial in term of energy efficiency: the brain can reduce the number of neurons for processing by using factorized representations, and switching only a part of assemblies requires a smaller number of synapses and plastic changes than overall control of the neural population. Our model confirmed that such efficient computational strategy actually works for semantic concepts learned from the text data in the real world.

### Relationships with hippocampal memory function

Further studies should clarify whether DSI can explain a wider range of hippocampal memory functions. A free recall of a given words is often used to evaluate memory functions in psychology (65, 66), and the underlying process has been modeled by Hopfield-type attractor neural network models (67–69). In these models, memory recall is interpreted as a transition across neural activity patterns embedded as attractors, and transition probabilities between these attractors are positively correlated with pattern similarity. Because DSI representations of two words become similar when the words share a semantic similarity, combining DSI with attractor network models likely enables us to investigate the relationship between semantic structures and the memory recall process. For example, such a model may explain the creation of false memory which is semantically similar to actually memorized items (65). Furthermore, the combination of recently learned associations in the hippocampus enables inference of novel relationships (70). An online extension of DSI may enable such inference based on one-shot memories.

### Alternative biological interpretations of DSI model

We interpreted conceptual units in our model as concept cells in HPC and EC because our model also produces place cells and grid cells. However, we can also interpret these units as “semantic features” (attributes) in a broad sense (43–45, 71, 72). A previous study has shown that distributed word representations composed of combinations of semantic features are useful for decoding words from fMRI activity patterns (71). It has been also shown that skip-gram representations support high-performance decoding of semantic information from fMRI data (73). In those experiments, semantic features are associated to brain-wide activity patterns, which implies that conceptual units in our model may also be related to brain areas other than HPC and EC. Especially, our model can be related to computations in two brain regions: anterior temporal lobe (ATL) and prefrontal cortex (PFC). It has been proposed that ATL processes semantic cognition and memory through interactions with multimodal sensory features across brain areas (43, 72). DSI model may be related to the computational mechanism of semantic features in ATL. As for PFC, fMRI studies have shown hexagonal modulations of neural activities in 2-D conceptual spaces as in EC (8–10). Furthermore, contributions of PFC to spatial navigation(74) and verbal analogical reasoning (75) have been suggested. Although concept-specific neural representations have not been found in PFC, other properties of DSI are consistent with those of PFC. These other possibilities should be investigated in future studies.

### Neural mechanism for natural language processing in the brain

Our model relates NLP, especially word embedding to conceptual representations in HPC and EC. Concept cells respond to words (both auditory inputs and texts) (25, 27), and the relationship between HPC and language processing have been experimentally found (76, 77). Regarding word embedding, a previous study showed that hippocampal theta oscillation codes semantic distances between words measured in word2vec (skip-gram) subspace (78). These experimental results support the contribution of HPC to language processing and word representations.

Recent studies have shown that representations in transformer-based models (79) such as GPT (80) achieve remarkable performance in linear fitting to neural recording during language processing (81, 82). Furthermore, it was recently shown that transformer-based models generate grid-like representations when applied to spatial learning (48). Similarly to our model, this finding implies the relationship between spatial and linguistic processing in the brain although concept-specific representations has not been found in transformer-based models. A major difference between our DSI model and transformer-based models is that DSI representations are basically fixed (static embedding) whereas transformer-based models flexibly create context-dependent representations (dynamic embedding). Computation of concepts obviously depends on the context; thus activities of concept cells are dependent on contexts such as memory contents and task demands (27). Therefore, our DSI model should be extended to process context-dependence, hopefully by combination with other models for learning context-dependent latent cognitive states (49, 83, 84).

### Relationships with disentangled visual representation learning

In our dimension reduction method, we introduced non-negativity, sparsity, decorrelation and regularization as constraints. Those constraints are also important for extraction of independent components (54, 85, 86), and it is known that imposing independence in latent spaces of deep generative models results in the emergence of disentangled representations for visual features (54, 64). As those disentangled visual representations explain single-cell activities in higher-order visual cortex (64), we similarly interpreted disentangled word representations in our model as concept cells. Concept cells respond to a specific concept, whereas population-level activity patterns represent abstract semantic structures (26). Such property is consistent with the factorized and distributed nature of disentangled representation vectors. Our model provides the view that concept cells emerge as disentangled representations for semantic concepts.

## Methods

### Mathematical relationship of SI and linear reinforcement learning

In this section, we show that SI corresponds to a value function of linear reinforcement learning (22–24) in the setting of spatial navigation.

In linear reinforcement learning, an agent aims to maximize “gain” instead of reward. Assuming a default policy

π^*d*^(*s*) (any policy is available; typically random walk in the case of exploration task), gain function is defined as

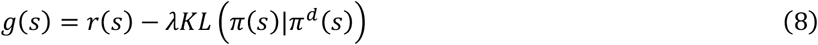

where *r*(*s*) is expected reward at the state *s* and *λKL* π(*s*)|π^*d*^(*s*) is the cost imposed on the difference between the current policy π(*s*) and the default policy π^*d*^(*s*) (*λ* is a relative weight of the cost). Then, previous works have shown that the optimal policy and corresponding value functions can be determined explicitly by solving linear equations (22–24). Here we consider an environment that consists of *N*_*N*_ nonterminal states and *N*_*T*_terminal states. We define two transition probability matrices under the default policy: P_*NT*_ is a *N*_*N*_ × *N*_*T*_matrix for transitions from non-terminal states to terminal states, and P_*NN*_ is a *N*_*N*_ × *N*_*N*_ matrix for transitions across non-terminal states. Furthermore, 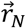 and 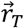 are vectors of rewards at non-terminal states and terminal states, respectively. In this condition, a vector of value functions under optimal policy 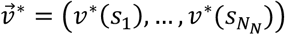 is obtained as

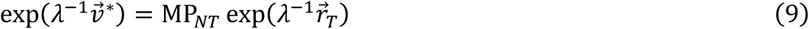

where 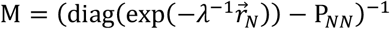 is DR (24).

To relate 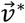 to SI, we consider a specific condition in which the environment consists of non-terminal states, and a virtual terminal state is attached to a goal state *s*_*G*_ arbitrarily chosen from those non-terminal states. When the agent gets to the goal state, it transits to the terminal state with a probability *p*_*NT*_. Furthermore, we assume that rewards at non-terminal states are uniformly negative and reward at the terminal state is positive so that the agent has to take a short path to goal to maximize reward. Specifically, we assume all elements of 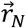 are *r*_−_, and *r*_*T*_= *r*_+_(*s*_*G*_) where *r*_−_ and *r*_+_(*s*_*G*_) is arbitrary negative and positive values, respectively. Then, we obtain

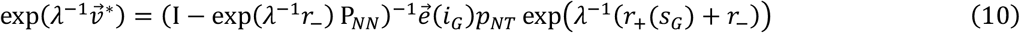

where *I* is an identity matrix, 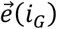 is a standard unit vector in which *i*_*G*_ -th element is 1, *i*_*G*_ is the index of the goal state. Because 0 < exp(*λ*^−1^*r*_−_) < 1, (*I* − exp(*λ*^−1^*r*_−_) P_*NN*_)^−1^ is equivalent to a successor representation matrix with a discount factor γ= exp(*λ*^−1^*r*_−_) (16, 18). Therefore, we obtain

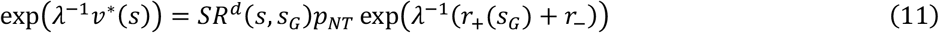

where *SR*^*d*^(*s, s*_*G*_) is SR under the default policy. By setting *r*_+_(*s*_*G*_) = −*r*_−_ − *λ* log *pp*_*NT*_− *λ* log *P*^*d*^(*s*_*G*_) (*P*^*d*^(*s*_*G*_) is a probability of visiting the state *s*_*G*_ under the default policy), we finally obtain

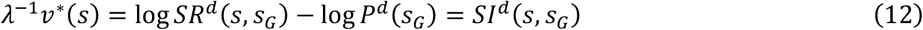

where *SI*^*d*^(*s, s*_*G*_) is SI under the default policy.

The transition probability matrix for non-terminal states P_*NN*_ slightly differs between settings with and without a goal because adding a transition to a terminal state changes transition probabilities from the goal state. Therefore, if SI is constructed through exploration of the environment without a goal (as in simulations in this paper), SI slightly deviates from the true value function. Rigorous matching is achieved by infinitesimal *p*_*NT*_(a small transition probability to the terminal state with a large positive reward) in that case.

### Calculation of SR

Throughout this study, we used a direct count method because we performed only offline processing of finite data. In a sequence of states {*s*_1_, …, *s*_*t*_, …, *s*_*T*_}, we recursively calculated exponential traces of past states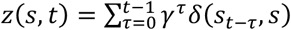 as

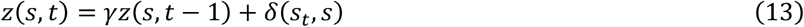

where *δ*(*i,j*) is Kronecker’s delta and *γ* is a discount factor. We calculated SR from state counts and coincidence counts as

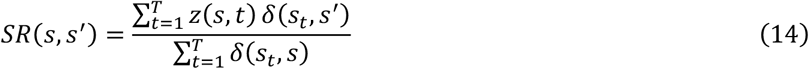

### Details of dimension reduction for DSI vectors

To obtain DSI-decorr, we iteratively updated *D*-dimensional vectors 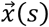 and 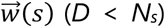 by Nesterov’s accelerated gradient descent method (87) to minimize the following objective function *J*_*decorr*_, rectifying all elements every iteration (*∀i, x*_*i*_ (*s*) ≥ 0, *w*_*i*_ (*s*) ≥ 0).

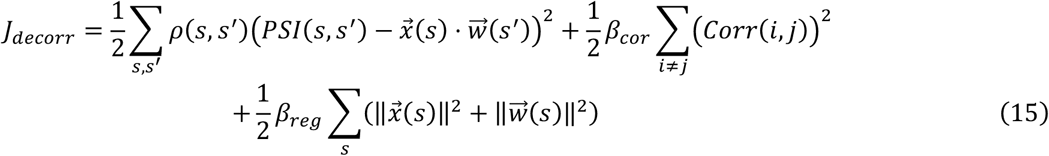

The first term of this objective function is weighted approximation error minimization, the second term works for decorrelation between dimensions, and the third term regularizes representation vectors. In this function, *ρ*(*s, s*^′^) is a weight for the square error

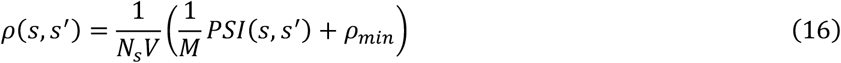

where *M* and *V* are mean and variance of PSI, respectively, and *ρ*_*min*_ is a small value to avoid zero-weight. *Corr*(*i, j*) is a correlation between two dimensions in 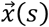

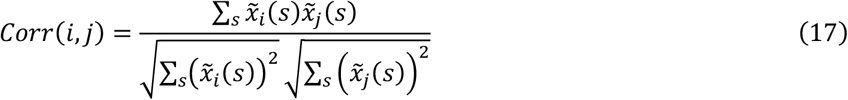

where 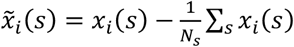.

Gradients of the objective functions for iterative updates are

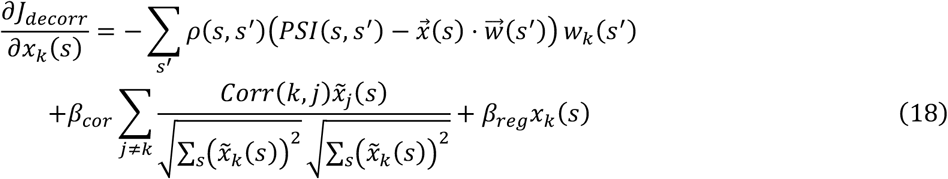

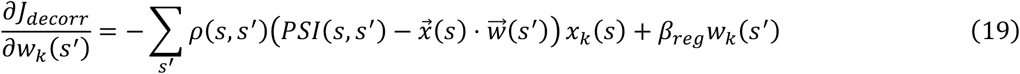

We note that we regarded mean and variance of *x*_*k*_(*s*) in the correlation 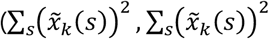 in Eq. 6) as constants in the calculation of these gradients. Practically, this heuristic did not affect the performance of decorrelation.

To obtain DSI-sparse, we performed the same procedure with DSI-decorr but we changed the objective function to

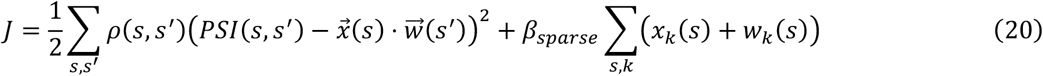

We note that the second term of this function is equivalent to a L-1 constraint because *x*_*k*_ (*s*) and *w*_*k*_ (*s*) are non-negative. Gradients are

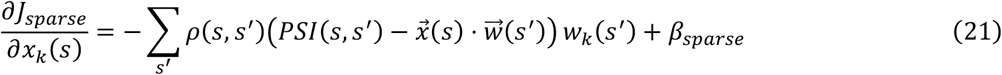

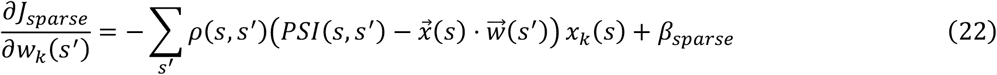

Throughout this paper, the learning rate was 0.05 and the number of iteration was 10000. Parameters were *β*_*cor*_ =1, *β*_*reg*_ = 0.001, *β*_*sparse*_=0.0001, and *ρρ*_*min*_ = 0.001.

### Learning DSI in 2-D spaces

As an environment, we assumed a square room (or interconnected separated rooms) tiled with 30 × 30 discrete states. In each simulation trial, an agent starts at one of those 900 states and transits to one of eight surrounding states each time except that transitions are limited at states along the boundary (the structure was not a torus). Transitions to surrounding states occur with an equal probability. We performed 500 simulation trials and obtained a sequence of 100,000 time steps in each trial. We calculated occurrence probabilities (*P*(*s*)) and a successor representation matrix (*SR*(*s,s*^′^)) of 900 states from those sequences, and calculated PSI and DSI (100-dimensional) as described in the Model section. The discount factor *γ* was set to 0.99.

We repeated this procedure five times using different random seeds which gave different representations each time, to report reproducibility or statistics of results.

### Gridness analysis

In analyses of spatial representations, we classified grid and non-grid cells based on gridness scores following the procedure in previous studies (37, 55–57). For each unit, we first determined the radius of a central peak in the spatial autocorrelation map following the criterion in previous study (56). We defined a circular area with outer radius *R* excluding the central peak (an annulus), centered on the origin of the spatial autocorrelation map. We calculated correlations between the original and rotated maps in the annulus. Gridness was defined as the difference between the lowest correlation at 60^°^ and 120^°^ and the highest correlation at 30^°^, 90^°^ and 150^°^. We repeated this calculation incrementing *R* from *R*_*min*_ to *R*_*max*_, and we determined gridness of the unit as a maximum value across all settings of *R*. For a simple 2-D room (the size was 30 × 30), *R*_*min*_ = 8 and *R*_*max*_ = 18. For “the context Φ” in spatial inference task (the size was 21 × 21), *R*_*min*_ = 6 and *R*_*max*_ = 14.

A unit was immediately classified as a non-grid cell when gridness did not exceed 0.3. For candidate grid cells (gridness>0.3), we performed a field-shuffling analysis (37, 57) to confirm that gridness is not obtained by chance. Briefly, we segmented spatial fields in a spatial representation map using a watershedding algorithm. We generated a shuffled map by randomly replacing each field to another position. We first replaced a peak bin, then bins around the peak were incrementally replaced, keeping the relative position to the peak bin. When the target bin had already been occupied by other fields, the nearest empty bin was used as the target bin instead. We created 100 shuffled maps and calculated gridness of shuffled maps to construct a null distribution. The unit was judged as a grid cell if gridness exceeds the 95 percentile of the null distribution.

### Path integration by DSI vectors

In the path integration task, we performed the estimation of states at each time step *s*_*t*_ from an initial state *s*_0_ and a sequence of movements {*a*_0_, *a*_1_, ⋯, *a*_*T*−1_} (*a*_*t*_ represents one of eight directional movements). To perform path integration, we initialized 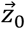 by a DSI representation vector 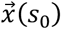, then we made an estimate of the next representation vector 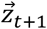 by linear transformation of the current representation vector 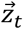 as

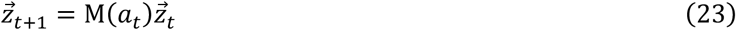

where M(*a*_*t*_) is movement-conditional recurrent weight matrix. We determined a position at each time step by searching a DSI representation vector 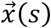 that has the minimum Euclidian distance with the estimated vector as

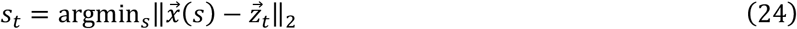

Before the estimation, we optimized the matrix M(*a*_*t*_) by minimizing prediction error 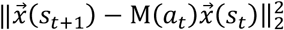 by stochastic gradient descent during random walk in the environment (20 simulation trials of 100,000 time steps).

### Goal-directed spatial navigation by DSI vectors

In each trial, we sampled a start location (state *s*_*init*_) and a goal location (state *s*_*G*_) such that the shortest path length is minimally 10, and an agent has to navigate between them. The rule of the state transition for navigation was as follows: suppose that the agent exists at a state *s*, and a set of neighboring states of *s* is *A*(*s*). Given the goal representation vector *w*(*s*_*G*_), a value of a neighboring state *s*_*next*_ *∈ A*(*s*) is estimated by *x*(*s*_*next*_) *⋅ w*(*s*_*G*_), and the agents transits to the state that has a maximum value.

Because of the relationship between value functions and DSI, this rule approximate value-based decision making for goal-directed navigation. However, because of approximation, the agent sometimes deviated from optimal paths. Specifically, the agent was sometimes trapped in local loops of a few adjacent states, which significantly impaired the performance. To avoid that, we heuristically introduced a “familiarity” variable *f*(*s*) to each state. This variable was initially set to zero and incremented by one each time the agent visits the state *s*. Then, value was evaluated by *x*(*s*_*next*_) *⋅ w*(*s*_*G*_)- *f*(*s*), which facilitates escape from local loops. Note that this strategy alone was not enough for appropriate goal-directed navigation (see results in Figure 7 for example).

In the evaluation, we terminated the simulation when the agent did not reach the goal in 100 time steps, and we considered the path length was 100 in the terminated trial.

### Learning DSI from text data

We used text data taken from English Wikipedia dump (enwiki-latest-pages-articles, 22-May-2020). We first generated text files from raw data using wikiextrator (https://github.com/attardi/wikiextractor). We tokenized texts by nltk Punkt sentence tokenizer, and randomly sampled 100,000 articles containing 1,000 tokens at minimum. We lowercased all characters and removed punctuation characters in the data. After that, we selected words that appeared more than 1,000 times in the data and substituted all other rare words by *<*unk*>* symbol. Finally, we obtained data that contains 124M tokens and 9376 words (i.e. 9376 states).

The procedure of calculating DSI was same as the case of 2-D spaces except that the discount factor *γ* was set to 0.9. The setting of other parameters was the same as the experiment of 2-D spaces. Using the preprocessed data, we repeated learning of DSI vectors five times using different random seeds (i.e. different initial values for optimization) which gave different representations each time, to report reproducibility or statistics of results.

### Comparison with other word embedding methods

We compared DSI with other word embedding models applied to the same text data under same dimensionality (300 dimensions). For skip-gram and continuous-bag-of-words (CBOW) (30, 31), we used implementation in Python Gensim library. For GLoVe (32), we used implementation by Pennington et al. (https://github.com/stanfordnlp/GLoVe). PPMI-SVD (33) and SR-SVD (18) were implemented by ourselves. The window size of coincidence count was set to 10 in all models, and other parameters followed default settings or parameters described in original papers. The discount rate of SR-SVD were same as SR used for calculation of DSI (γ= 0.9). For BERT, we took representations in a pretrained model (bert-base-uncased in Hugging Face transformers) (60, 61), thus training data and dimensionality (768 dimensions) were different from our setting.

### Quantitative evaluation of conceptual specificity

Conceptual specificity of each unit was evaluated using WordNet database (59). In WordNet, a word belongs to several synsets (sets of cognitive synonyms), and semantic similarity of two synsets can be evaluated from the shortest path length between them in the WordNet structure (we used path similarity function in nltk library). We defined similarity of two words as the highest similarity among all combinations of synsets of those words. We calculated mean similarity of all combinations of TOP-10 words (ten words that highly activated the unit; Figure 4A) that are available in WordNet. We evaluated only units which had at least five TOP-10 words available in WordNet. Furthermore, we randomly generated 1,000 pairs of words available in WordNet, and generated a null distribution of similarity between words. We defined a significance threshold of similarity as a 95 percentile of the null distribution, and a unit was classified as a significantly concept-specific unit if mean similarity of TOP-10 words exceeded the threshold. Furthermore, we quantitatively defined a conceptual specificity of each unit as

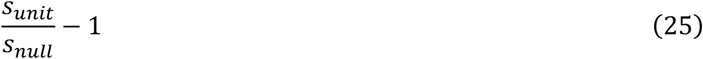

where *s*_*unit*_ is mean similarity of TOP-10 words and *s*_*null*_ is the mean of the null distribution. This quantity becomes zero if similarity between TOP-10 words is not different from random pairs, and becomes positive if TOP-10 words are semantically similar. This conceptual specificity was averaged over all evaluated units.

Of note, we found several non-significant units exhibit conceptual specificity according to manual inspection (see Supplementary Figure 3). This is probably because of the limitation of knowledge covered by WordNet. Therefore, we suppose that the current evaluation method tends to underestimate the number of concept-specific units. However, the comparison across models was fair because we used the same procedure and criteria for all models.

### Evaluation of the semantic structure of DSI vectors

To analyze the semantic structure in representation vectors, we considered ten semantic categories (“mammal”, “building”, “vehicle”, “food”, “clothing”, “body part”, “computer”, “feeling”, “creator”, “relative”) and chose ten words for each category as follows.

“mammal”: dog, cat, bull, bear, elephant, fox, horse, lion, rat, tiger

“building”: building, church, theater, hall, school, hotel, house, library, mansion, palace

“vehicle”: vehicle, ambulance, bike, aircraft, bus, car, helicopter, locomotive, rocket, ship

“food”: food, bread, cheese, candy, rice, beer, coffee, milk, tea, wine

“clothing”: clothing, shirt, hat, belt, costume, dress, wear, cap, coat, crown

“body part”: arm, bone, lung, ear, eye, finger, foot, hair, kidney, leg

“computer”: computer, software, internet, code, access, server, website, pc, node, portal

“feeling”: feeling, anger, anxiety, comfort, confusion, emotion, enthusiasm, happiness, joy, love

“creator”: creator, architect, artist, composer, designer, producer, filmmaker, photographer, musician, painter

“relative”: husband, brother, aunt, cousin, daughter, father, grandfather, grandmother, mother, sister

We took DSI representation vectors for these 100 words and computed dissimilarity (1-Pearson’s correlation coefficient). MDS was performed using the same dissimilarity metric.

### Analogical inference of words

In Mikolov’s dataset, many sets of four words are given. There is a relationship “WORD1 is to WORD2 as WORD3 is to WORD4” (i.e. king is to queen as brother is to sister). Then, an expected relationship in the vector space is WORD2-WORD1=WORD4-WORD3. In this study, we performed inference of WORD4 by WORD3+WORD2-WORD1. We regarded an inference was correct if the actual vector of WORD4 had the largest cosine similarity to the inferred vector among all word representation vectors (except those for WORD1, WORD2, and WORD3). If the number of words is 10,000, a chance level of the correct answer rate is 0.01%. Therefore, the performance shown in this study (more than 50%) is far above the chance level.

When we performed analogical inference by calculation of the limited number of dimensions (partial recombination), we first calculated 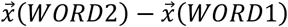 and then identified the *n* largest and the *n* smallest elements in that vector. We summed those 2*n* out of 300 elements to 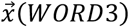 (in other words, other elements of 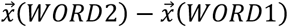 were substituted by zero before summation). Correctness of the inference was evaluated by the same way with the original task.

### Analogical inference of spatial contexts

We considered a 2-D space tiled with 21×21 (441) states and four spatial contexts (barrier layouts) A, B, Φ, A+B (Supplementary Figure 6). We define separated states in contexts A, B, Φ, A+B as 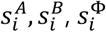 and 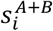 where *i* is a positional index which indicates a same position in all contexts (*i* = 1,2, …, 441). We constructed representation vectors for contexts A, B, Φ through direct experiences, then we created 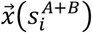 and 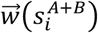 as

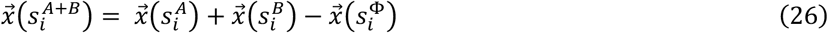

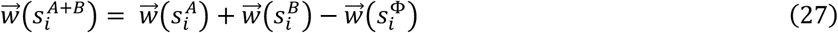

We performed spatial navigation in a given context using one of four representations 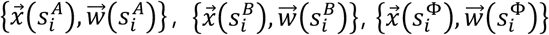 and 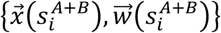 for corresponding positions, following the vector-based decision making rule described in the previous section (“Goal-directed spatial navigation by DSI vectors”).

To learn representations in contexts A, B, and Φ, we sampled sequences of 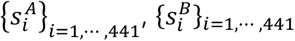 ^, and^ 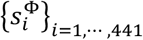 as in the previous section (“Learning DSI in 2-D spaces”). However, we performed random walkin the 2-D space while switching between context A, B, and Φ, that is, state transitions to the same position in other contexts occurred every 5,000 time steps (transition between 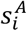 and 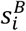, and 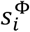). It means that we assumed that the setting of barriers can change during the experience. This transition was introduced to associate the same position in different contexts. We performed 500 simulation trials and obtained a sequence of 100,000 time steps in each trial. From sampled sequences, we calculated SR for all combinations of 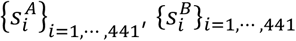 and 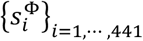 and calculated 100-dimensional DSI vectors for 1,324 (441×3) states by simultaneous compression of all states (dimension reduction of a 1,324×1,324 PSI matrix). The discount factor *γ* was set to 0.99.

When we performed the inference by calculation of the limited number of dimensions, we first identified the *n* units (dimensions) that had the largest representational distances between the context B and Φ. The representational distance of the unit *k* (*k*-th dimension of vectors *x*_*k*_ (*s*)) was defined as

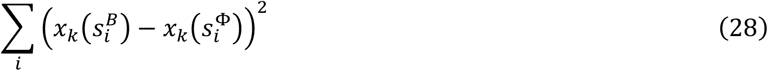

We calculated 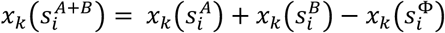 only for those *n* dimensions, and we set 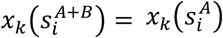 in other dimensions.

### Statistical analysis

To ensure reproducibility, we showed results obtained by five repetitions of simulations with different random seeds (different initial values for learning and simulations) in Figure 4, Figure 6, Figure 7, Supplementary Figure 7 and 8. We applied two-sided t-tests to those five samples assuming normality of data, and detailed values of t-statistic and p-values are summarized in Supplementary Table 1, 2, and 3. Significance thresholds were modified by Bonferroni correction.

## Code and data availability

Python codes for simulations and analyses are available at https://github.com/TatsuyaHaga/DSI_codes. We also used external codes at https://github.com/stanfordnlp/GLoVe (GLoVe) and https://github.com/attardi/wikiextractor (wikiextrator). Text data were taken from https://dumps.wikimedia.org/enwiki/latest/ but currently the version 22-May-2020 is not available. We share the version we used upon request. WS353 dataset is available at http://alfonseca.org/eng/research/wordsim353.html. Mikolov’s dataset is available at https://aclweb.org/aclwiki/Google_analogy_test_set_(State_of_the_art).

## Acknowledgments

We thank Ryo Yoshida for supporting the analysis of representations in BERT model. We also thank Robert M Mok for fruitful discussion. This research was partially supported by JSPS KAKENHI Grant Numbers 21K15611, 19H04994, 18H05213, and JST PRESTO Grant Number JPMJPR21C2.

## Author contributions

T.H.: conceptualization, formal analysis, funding acquisition, investigation, methodology, software, visualization, writing-original draft, writing-review & editing. Y.O.: conceptualization, funding acquisition, supervision, writing-review & editing. T.F.: conceptualization, funding acquisition, resources, supervision, writing-review & editing.

