## Supplementary materials for "A unified neural representation model for spatial and semantic computations"

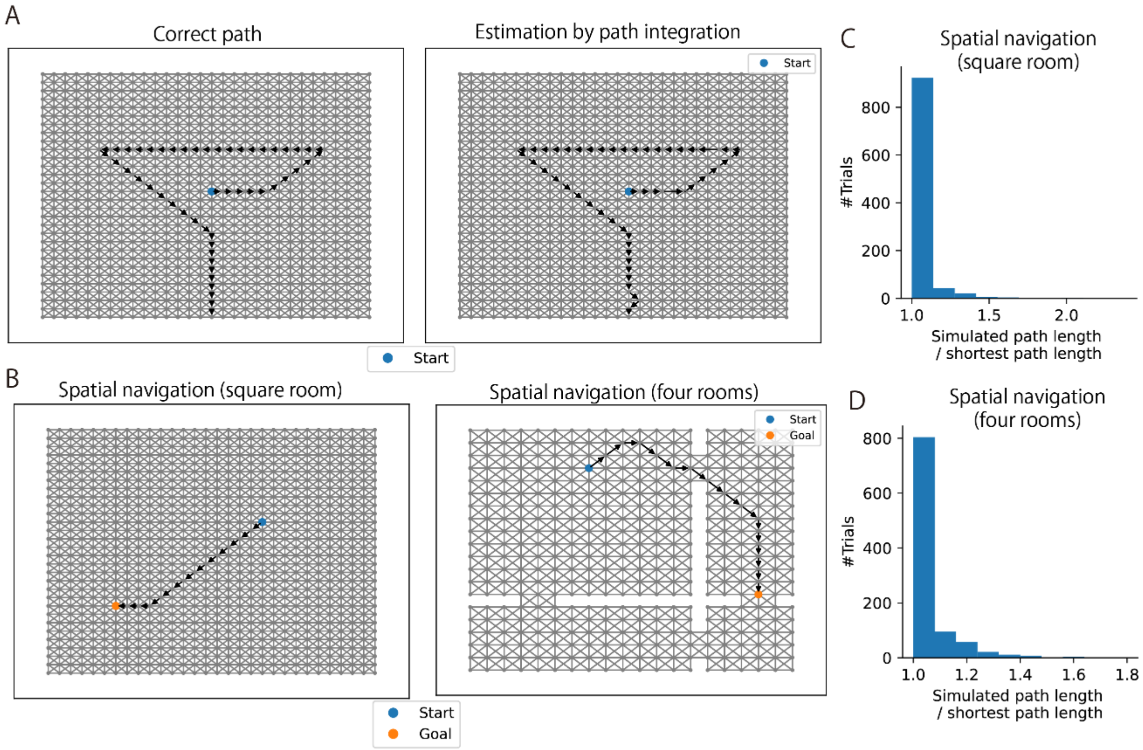


Supplementary Figure 1: Spatial navigation using DSI representation vectors (DSI-sparse). (A) Path integration using DSI model. (Left) Actual path. (Right) Path estimated from DSI vectors updated by movement information. (B) Example spatial paths obtained by DSI-based navigation. (C) A histogram of path lengths in 1,000 trials of spatial navigation in the square-room environment. Note that a start and a goal were randomly determined in each trial, and we normalized a simulated path length by the shortest path length between the start and the goal. (D) A histogram of path lengths in the four-room environment.

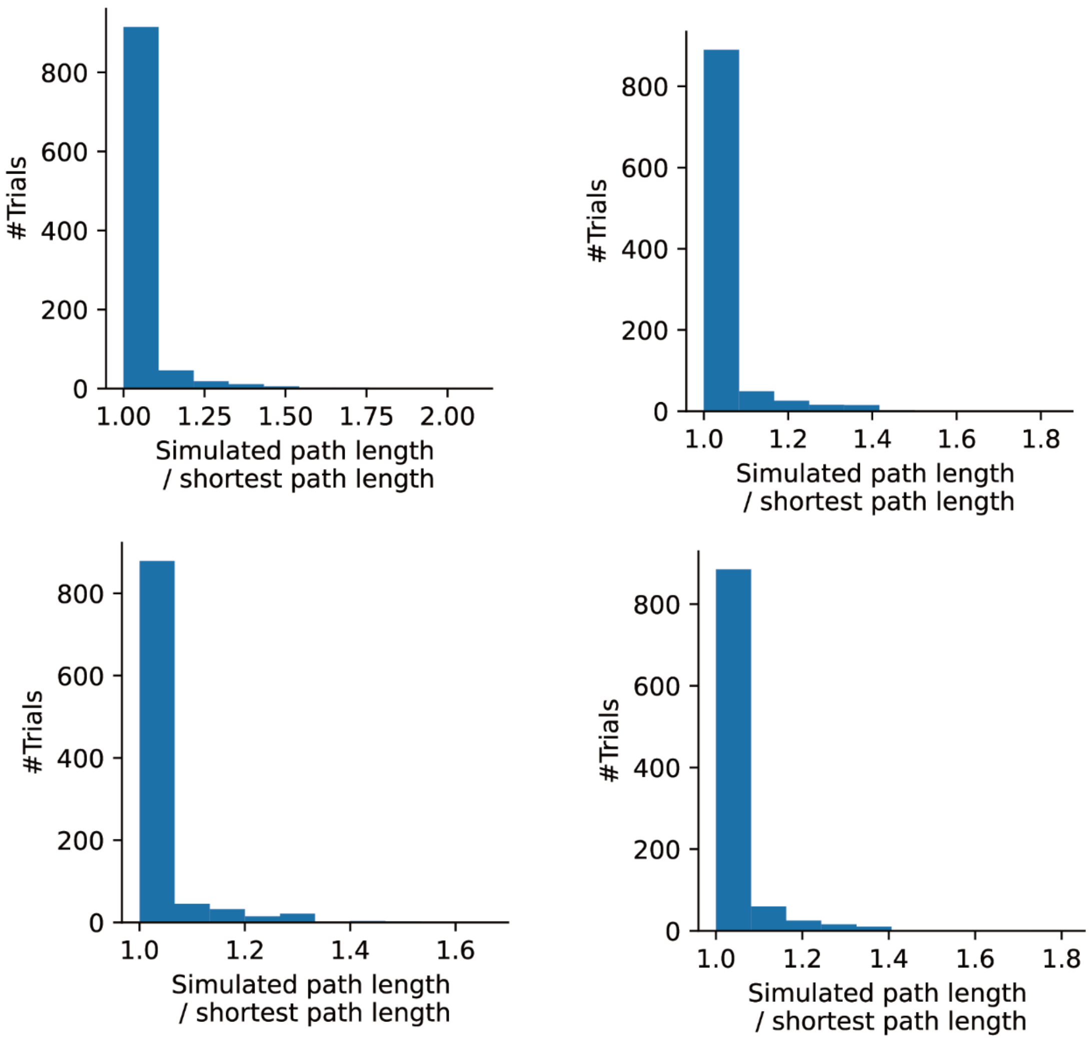


Supplementary Figure 2: Histograms of path lengths in 1,000 trials of spatial navigation in the square-room environment obtained by 4 simulations with different random seeds. Note that a start and a goal were randomly determined in each trial, and we normalized a simulated path length by the shortest path length between the start and the goal.

| Unit 1 | Unit 2 | Unit 3 | Unit 4 | Unit 5 |
|---|---|---|---|---|
| faced | transit | humid | universities | sheets |
| overcome | commuter | winters | colleges | olivia |
| facing | trains | summers | uc | compact |
| due | bus | climate | usc | senators |
| owing | buses | mild | consortium | antony |
| citing | subway | precipitation | institutes | proposes |
| experiencing | rail | warm | carnegie | atlas |
| cope | passenger | temperatures | affiliated | spaces |
| amid | metro | cool | alumni | geometry |
| suffered | passengers | cold | selective | rough |

| Unit 6 | Unit 7 | Unit 8 | Unit 9 | Unit 10 |
|---|---|---|---|---|
| designers | rebounds | nbc | eleventh | nhl |
| scientists | assists | aired | tenth | confirmation |
| experts | averaged | abc | ninth | hockey |
| professionals | steals | airing | seventh | maple |
| consumers | nba | cbs | eighth | roller |
| researchers | averaging | syndicated | twelfth | judiciary |
| filmmakers | partition | programming | 7th | flames |
| lawyers | points | broadcast | 8th | devils |
| composers | prussia | channel | 10th | batch |
| scholars | mvp | espn | 11th | senators |

| Unit 11 | Unit 12 | Unit 13 | Unit 14 | Unit 15 |
|---|---|---|---|---|
| lowest | valve | archive | hatch | rotten |
| ranked | wheel | archives | deaf | tomatoes |
| highest | steering | manuscripts | robot | metacritic |
| ranks | rotating | collections | suffrage | aggregate |
| literacy | wheels | collection | egg | approval |
| demographic | cylinder | manuscript | seymour | reviews |
| rising | trigger | library | salem | grossed |
| prevalence | shaft | catalogue | wwe | rating |
| ratings | pump | valuable | monte | byron |
| rank | gear | documents | resist | apparatus |

Supplementary Figure 3: A part of word representations formed by DSI-decorr. Ten words that gave the highest activation (TOP-10 words) are shown for units 1 - 15. Those classified as concept-specific units in our analysis are boxed with bold lines.

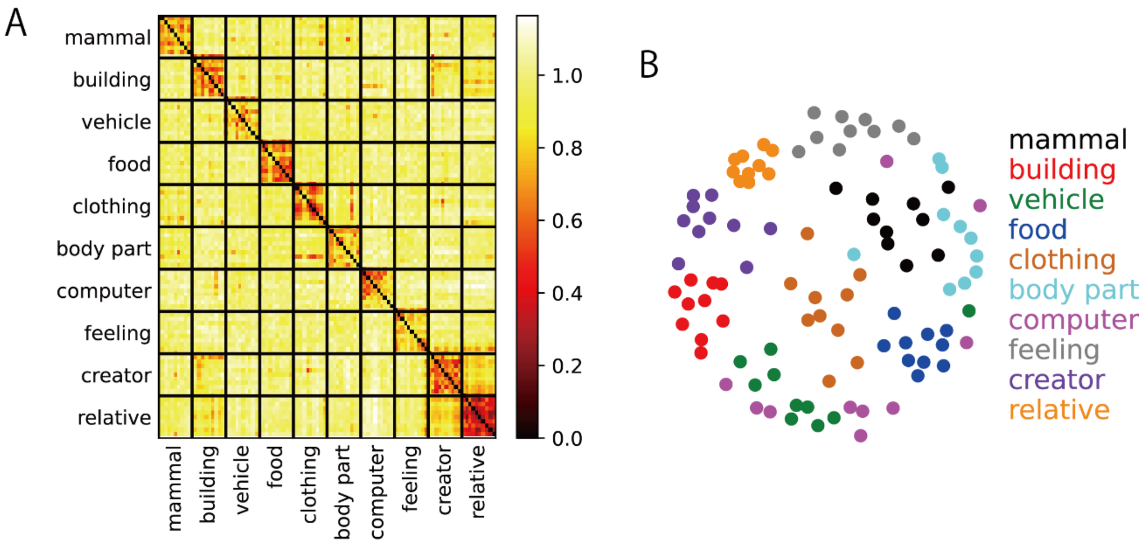


Supplementary Figure 4: DSI representations capture the semantic structure of words at a population level (DSI-sparse). (A) Dissimilarity matrix between DSI representation vectors for 100 words in 10 semantic categories. We selected 10 words in each category. We used same dissimilarity metric with Reber et al. (2019) (1 - Pearson's correlation coefficient). (B) Visualization of the representational structure of DSI using MDS based on the dissimilarity matrix. Each color corresponds to a semantic category.

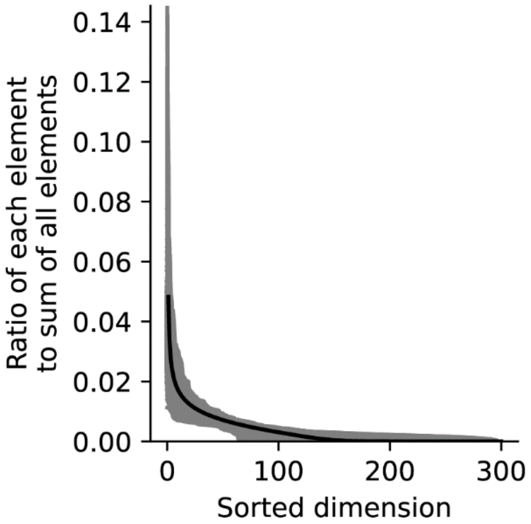


Supplementary Figure 5: Non-sparsity of word representations. Gray lines show ratio of each element to sum of all elements in each word representation vector, and the black line is the average of them. Elements were sorted in descending order.

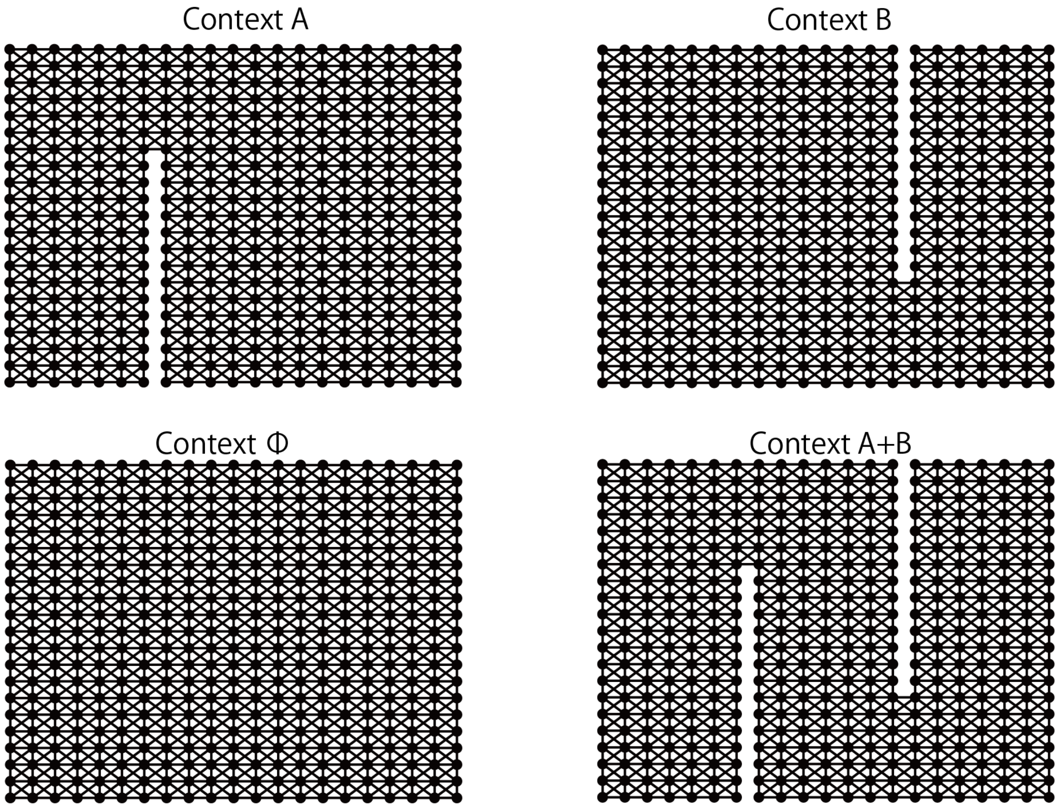


Supplementary Figure 6: Structures of state transition graphs used in inference task of spatial contexts.

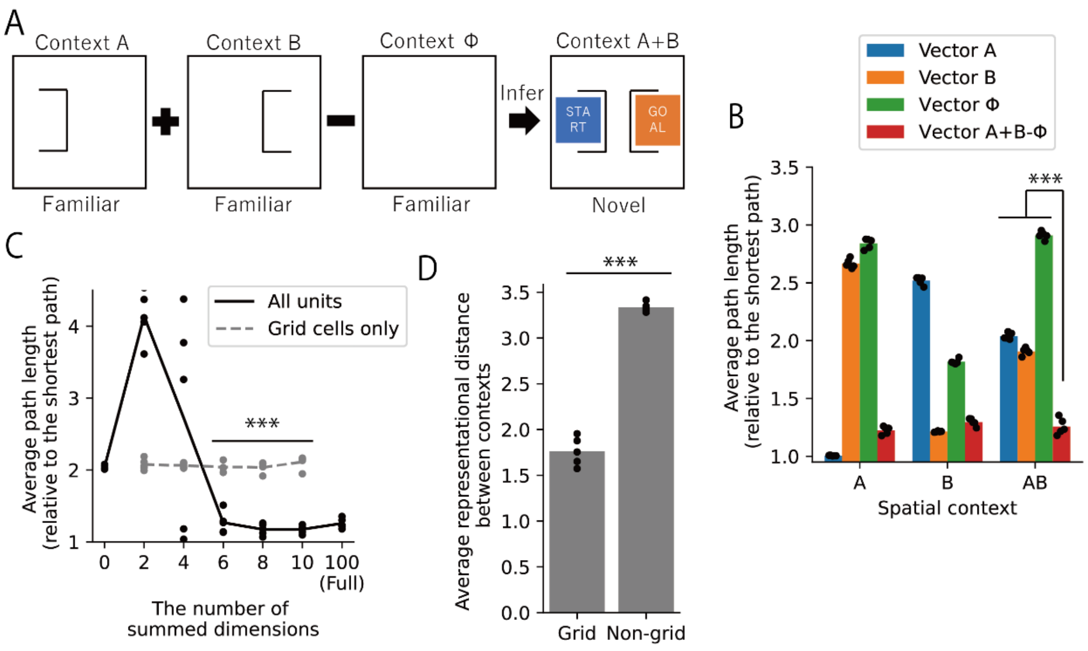


Supplementary Figure 7: Composite spatial representations enable navigation in a novel spatial context. (A) We constructed representation vectors for a novel context A+B by arithmetic composition of DSI representation vectors for three familiar contexts A, B, and Φ. The start and the goal in each navigation trial were randomly positioned in the colored area. (B) Average path lengths in 1,000 trials of spatial navigation under various settings of representation vectors and contexts. Note that we normalized a path length by the shortest path length between the start and the goal in each trial. Dots indicate 5 simulations with different random seeds (different initial values for learning and simulations); bars indicate means of those 5 simulations. (C) Average path lengths by the spatial navigation using the composite representation vectors in which we summed only the limited number of dimensions. Dots indicate 5 simulations with different random seeds (different initial values for learning and simulations); bars indicate means of those 5 simulations. 2-sample t-tests were performed between the condition in which we calculated only grid cells and the condition in which we calculated all units. (D) Average representational distances between the context B and Φ across 100 units (dimensions). Dots indicate 5 simulations with different random seeds (different initial values for learning and simulations); bars indicate means of those 5 simulations. ***$P<0.001$. All statistical tests were two-sided t-tests and significance thresholds were modified by Bonferroni correction. Details of statistical analyses are shown in Supplementary Table 3.

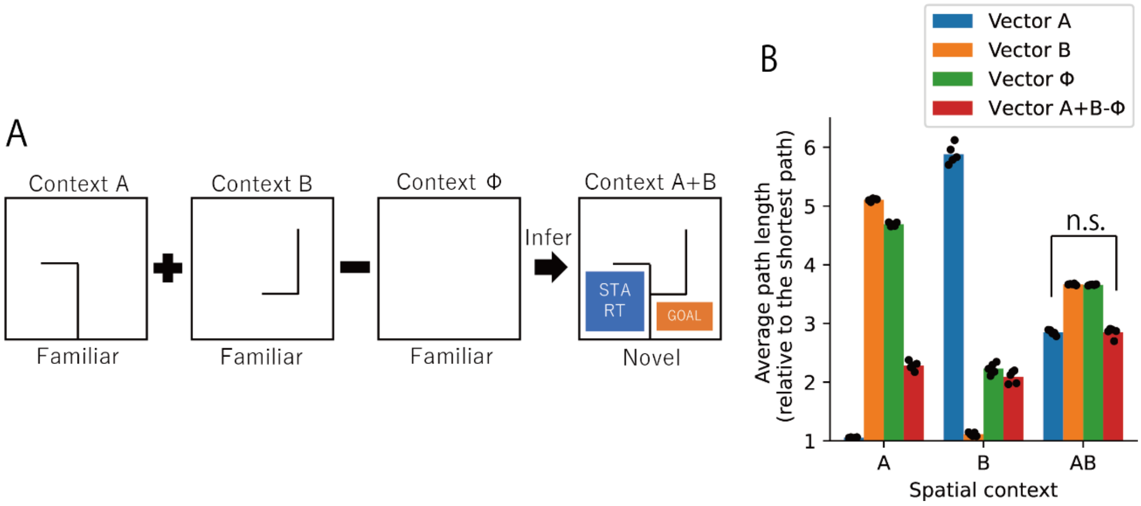


Supplementary Figure 8: An example in which composite spatial representations did not enable navigation in a novel spatial context. (A) We constructed representation vectors for a novel context A+B by arithmetic composition of DSI representation vectors for three familiar contexts A, B, and Φ. The start and the goal in each navigation trial were randomly positioned in the colored area. (B) Average path lengths in 1,000 trials of spatial navigation under various settings of representation vectors and contexts. Note that we normalized a path length by the shortest path length between the start and the goal in each trial. Dots indicate 5 simulations with different random seeds (different initial values for learning and simulations); bars indicate means of those 5 simulations. n.s., not significant. All statistical tests were two-sided t-tests and significance thresholds were modified by Bonferroni correction. Details of statistical analyses are shown in Supplementary Table 3.

Supplementary Table 1: Details of statistical analyses in Figure 3.

| Comparison | Statistical test | Statistic | p-value |
|---|---|---|---|
| Figure 3B | | | |
| DSI (decorr) vs DSI (sparse) | Two-sided 2-sample t-test | t(8)=1.2 | p=0.27 |
| DSI (decorr) vs CBOW | Two-sided 1-sample t-test | t(4)=1.2 | p=0.91 |
| DSI (decorr) vs DSI (non-neg. OFF) | Two-sided 2-sample t-test | t(8)=12.5 | $p=1.5\times10^{-6}$ |
| DSI (decorr) vs Skip-gram | Two-sided 1-sample t-test | t(4)=12.3 | $p=2.5\times10^{-4}$ |
| DSI (decorr) vs GLoVe | Two-sided 1-sample t-test | t(4)=13.8 | $p=1.6\times10^{-4}$ |
| DSI (decorr) vs PPMI-SVD | Two-sided 1-sample t-test | t(4)=8.4 | p=0.0011 |
| DSI (decorr) vs SR-SVD | Two-sided 1-sample t-test | t(4)=8.0 | p=0.0013 |
| DSI (decorr) vs BERT | Two-sided 1-sample t-test | t(4)=11.1 | $p=3.8\times10^{-4}$ |
| Figure 3C | | | |
| DSI (decorr) vs DSI (sparse) | Two-sided 2-sample t-test | t(8)=1.4 | p=0.21 |
| DSI (decorr) vs CBOW | Two-sided 1-sample t-test | t(4)=0.9 | p=0.41 |
| DSI (decorr) vs DSI (non-neg. OFF) | Two-sided 2-sample t-test | t(8)=17.6 | $p=1.1\times10^{-7}$ |
| DSI (decorr) vs Skip-gram | Two-sided 1-sample t-test | t(4)=17.5 | $p=6.2\times10^{-5}$ |
| DSI (decorr) vs GLoVe | Two-sided 1-sample t-test | t(4)=15.7 | $p=9.5\times10^{-5}$ |
| DSI (decorr) vs PPMI-SVD | Two-sided 1-sample t-test | t(4)=9.1 | $p=8.2\times10^{-4}$ |
| DSI (decorr) vs SR-SVD | Two-sided 1-sample t-test | t(4)=8.3 | p=0.0011 |
| DSI (decorr) vs BERT | Two-sided 1-sample t-test | t(4)=10.7 | $p=4.3\times10^{-4}$ |

Supplementary Table 2: Details of statistical analyses in Figure 5D.

| Number of calculated dimensions | n=2 | n=4 | n=6 | n=8 | n=10 |
|---|---|---|---|---|---|
| DSI (decorr) vs DSI (sparse), Two-sided 2-sample t-test | t(8)=1.9, p=0.10 | t(8)=1.3, p=0.22 | t(8)=1.6, p=0.14 | t(8)=1.1, p=0.21 | t(8)=1.2, p=0.28 |
| DSI (decorr) vs DSI (non-neg. OFF), Two-sided 2-sample t-test | t(8)=22.9, $p=1.4 \times 10^{-8}$ | t(8)=18.1, $p=8.9 \times 10^{-8}$ | t(8)=17.4, $p=1.2 \times 10^{-7}$ | t(8)=16.8, $p=1.6 \times 10^{-7}$ | t(8)=12.3, $p=1{,}8 \times 10^{-6}$ |
| DSI (decorr) vs CBOW, Two-sided 1-sample t-test | t(4)=45.2, $p=1.4 \times 10^{-6}$ | t(4)=32.5, $p=5.4 \times 10^{-6}$ | t(4)=34.1, $p=4.4 \times 10^{-6}$ | t(4)=34.7 $p=4.1 \times 10^{-6}$ | t(4)=28.8, $p=8.7 \times 10^{-6}$ |
| DSI (decorr) vs Skip-gram, Two-sided 1-sample t-test | t(4)=40.1, $p=2.3 \times 10^{-6}$ | t(4)=29.7, $p=7.6 \times 10^{-6}$ | t(4)=32.5, $p=5.3 \times 10^{-6}$ | t(4)=33.6, $p=4.7 \times 10^{-6}$ | t(4)=28.5, $p=9.0 \times 10^{-6}$ |
| DSI (decorr) vs GLoVe, Two-sided 1-sample t-test | t(4)=39.8, $p=2.4 \times 10^{-6}$ | t(4)=30.7, $p=6.7 \times 10^{-6}$ | t(4)=33.9, $p=4.5 \times 10^{-6}$ | t(4)=37.5, $p=3.0 \times 10^{-6}$ | t(4)=32.5, $p=5.3 \times 10^{-6}$ |
| DSI (decorr) vs PPMI-SVD, Two-sided 1-sample t-test | t(4)=41.7, $p=2.0 \times 10^{-6}$ | t(4)=28.6, $p=8.9 \times 10^{-6}$ | t(4)=28.8, $p=8.7 \times 10^{-6}$ | t(4)=30.1, $p=7.2 \times 10^{-6}$ | t(4)=25.1, $p=1.5 \times 10^{-4}$ |
| DSI (decorr) vs SR-SVD, Two-sided 1-sample t-test | t(4)=59.0, $p=5.0 \times 10^{-7}$ | t(4)=46.3, $p=1.3 \times 10^{-6}$ | t(4)=53.4, $p=7.4 \times 10^{-7}$ | t(4)=61.7, $p=4.1 \times 10^{-7}$ | t(4)=56.9, $p=5.7 \times 10^{-7}$ |

Supplementary Table 3: Details of statistical analyses in Figure 6, Supplementary Figure 7 and 8. p=0 indicates a value smaller than the minimum of the floating-point number.

| Comparison | Statistical test | Statistic | p-value |
|---|---|---|---|
| Figure 6C | | | |
| Vector A+B-Φ vs Vector A | Two-sided 2-sample t-test | t(8)=52.1 | p=0 |
| Vector A+B-Φ vs Vector B | Two-sided 2-sample t-test | t(8)=90.3 | p=0 |
| Vector A+B-Φ vs VectorΦ | Two-sided 2-sample t-test | t(8)=96.8 | p=0 |
| Figure 6E | | | |
| All units vs Grid cells only, n=2 | Two-sided 2-sample t-test | t(8)=2.4 | p=0.043 |
| All units vs Grid cells only, n=4 | Two-sided 2-sample t-test | t(8)=46.7 | p=0 |
| All units vs Grid cells only, n=6 | Two-sided 2-sample t-test | t(8)=20.5 | p=0 |
| All units vs Grid cells only, n=8 | Two-sided 2-sample t-test | t(8)=22.1 | p=0 |
| All units vs Grid cells only, n=10 | Two-sided 2-sample t-test | t(8)=28.3 | p=0 |
| Figure 6H | | | |
| Grid cells vs Non-grid cells | Two-sided 2-sample t-test | t(8)=24.5 | p=0 |
| Supplementary Figure 7B | | | |
| Vector A+B-Φ vs Vector A | Two-sided 2-sample t-test | t(8)=23.1 | p=0 |
| Vector A+B-Φ vs Vector B | Two-sided 2-sample t-test | t(8)=18.9 | p=0 |
| Vector A+B-Φ vs VectorΦ | Two-sided 2-sample t-test | t(8)=47.3 | p=0 |
| Supplementary Figure 7C | | | |
| All units vs Grid cells only, n=2 | Two-sided 2-sample t-test | t(8)=12.8 | $p=1\times10^{-6}$ |
| All units vs Grid cells only, n=4 | Two-sided 2-sample t-test | t(8)=0.97 | p=0.36 |
| All units vs Grid cells only, n=6 | Two-sided 2-sample t-test | t(8)=10.4 | $p=6\times10^{-6}$ |
| All units vs Grid cells only, n=8 | Two-sided 2-sample t-test | t(8)=17.9 | p=0 |
| All units vs Grid cells only, n=10 | Two-sided 2-sample t-test | t(8)=19.4 | p=0 |
| Supplementary Figure 7D | | | |
| Grid cells vs Non-grid cells | Two-sided 2-sample t-test | t(8)=21.3 | p=0 |
| Supplementary Figure 8B | | | |
| Vector A+B-Φ vs Vector A | Two-sided 2-sample t-test | t(8)=0.03 | p=0.98 |
